# Excitable dynamics of NREM sleep: a unifying model for neocortex and hippocampus

**DOI:** 10.1101/312587

**Authors:** Daniel Levenstein, György Buzsáki, John Rinzel

## Abstract

During non-rapid eye movement (NREM) sleep, the neocortex and hippocampus alternate between periods of neuronal spiking and inactivity. By directly comparing experimental observations with a mean field model of an adapting, recurrent neuronal population, we find that the neocortical alternations reflect a dynamical regime in which a stable active state is interrupted by transient inactive states (slow waves) while the hippocampal alternations reflect a stable inactive state interrupted by transient active states (sharp waves). We propose that during NREM sleep, hippocampal and neocortical populations are excitable: each in a stable state from which internal fluctuations or external perturbation can evoke the stereotyped population events that mediate NREM functions.

---

Sleep function relies on internally-generated dynamics in neuronal populations. In the neocortex, non-rapid eye movement (NREM) sleep is dominated by a “slow oscillation”^1^: alternations between periods of spiking (UP states) and periods of hyperpolarization (DOWN states) that correspond to large “slow waves” (or “delta waves”) in the local field potential (LFP)^2,3^ (Figure 1A,B, Supplemental Figure 1). In the hippocampus, NREM sleep is dominated by sharp wave-ripple dynamics: periods of spiking (SWRs) separated by periods of relative inactivity (inter-SWRs)^4^ (Figure 1E,F). Slow waves and SWRs are bidirectionally and weakly coupled, in that each is more likely following the other^5-8^. The functional importance of these dynamics is well established: both slow waves and SWRs have been observed to perform homeostatic maintenance of the local synaptic network in the two regions^9-11^, and their temporal coupling has been found to support the consolidation of recently learned memories^12-14^. However, it’s unclear how the state of neuronal populations in the two regions promotes the generation of their respective dynamics, or how population state supports the propagation of neural activity between structures.

**Figure 1:**
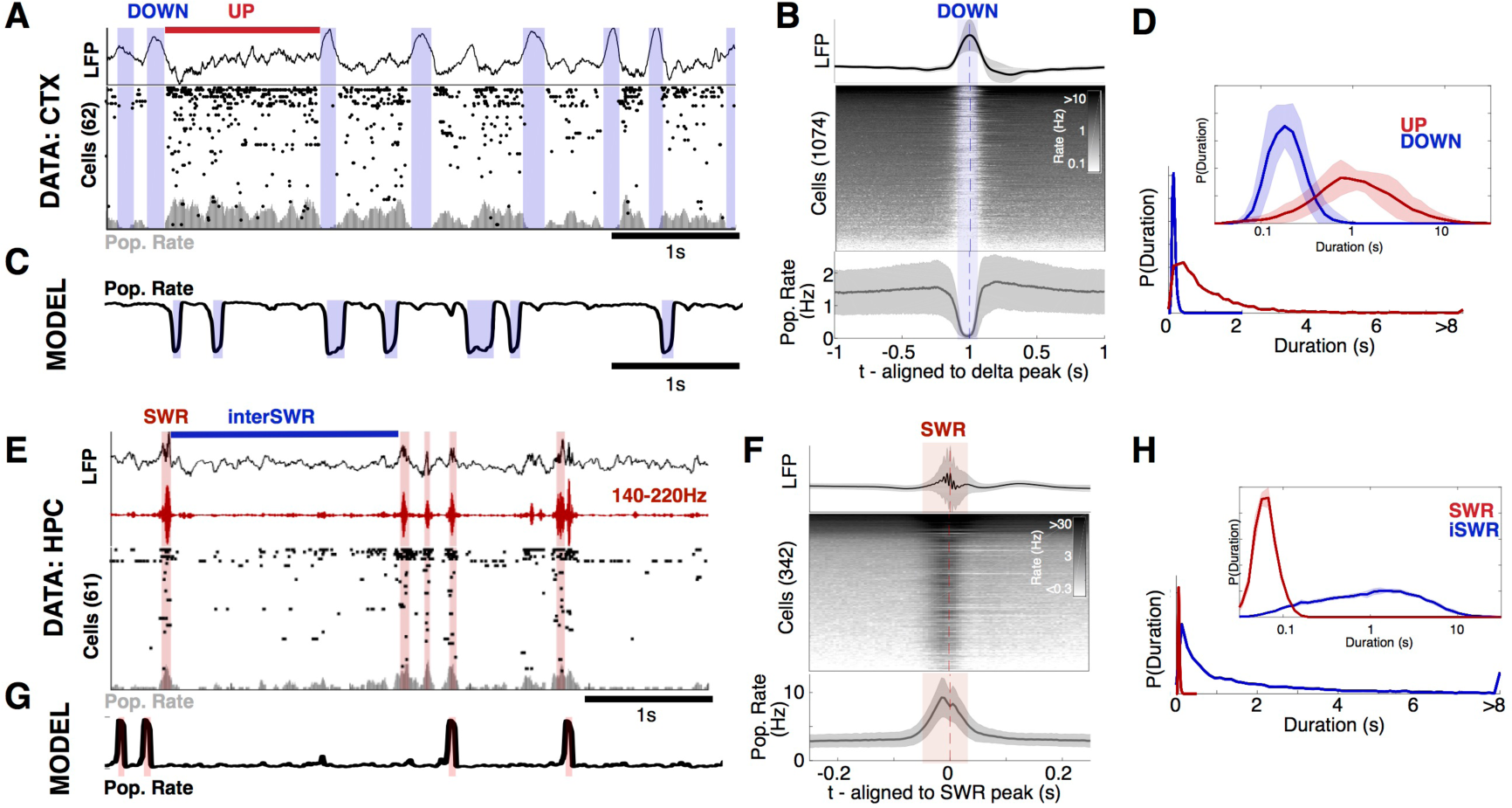
Neocortical UP/DOWN and hippocampal SWR dynamics during NREM sleep. **A:** A sample of data from rat mPFC during NREM sleep. Data were collected using high-density silicon probes (Watson et al 2016). LFP and spike times from cortical neurons were extracted as reported previously (see Methods). Neocortical slow waves are coincident with population-wide non-spiking **DOWN** states, which alternate with UP states of longer duration. **B:** Peri-event time histogram aligned to delta peaks for the LFP (top), all recorded cells (middle), and population rate (bottom). Mean +/ standard deviation over all recordings shown. C: Simulation of the model (eqns 1-2). Parameters determined by matching in vivo and simulated UP/DOWN state dwell times, as described in section: *Neocortex* is *in an Excitableup regime during NREM sleep.* **D:** UP/DOWN state dwell time distribution (bottom) in linear (example recording) and logarithmic scale (mean+/-std over all recordings). **E: A** sample of data from rat CA1 (HPC) during NREM sleep (Grosmark and Buzsaki 2016). Detected sharp wave-ripples (SWR) state indicated with red. **F:** Peri-event time histogram aligned to SWR peaks for the LFP (top), all recorded cells (middle), and population rate (bottom). Mean +/- standard deviation over all recordings shown. G: Simulated *r-a* model with the best-matching parameters, as described in section: *Hippocampus* is *in an ExcitableoowN regime during NREM sleep.* **H:** SWR and inter-SWR duration distributions in linear (example recording) and logarithmic scale (mean+/-std over all recordings).

To study the state of hippocampal and neocortical populations during NREM sleep, we used an idealized model of an adapting recurrent neuronal population (Figure 1C,G). Models with recurrence and adaptation have been directly matched to neocortical UP/DOWN alternations during anesthesia and in slice preparations^15-17^. These studies found that the UP/DOWN alternations in slice are adaptation-mediated oscillations^16^, while those in the anesthetized animal reflect noise-induced switches between bistable states^15^. However neuronal dynamics during NREM sleep in naturally sleeping rats^18^ are distinct from those seen in anesthesia/slice^19^. With our adapting recurrent population model we are able to describe how effective physiological parameters determine the properties of alternation dynamics in a neuronal population. This framework allowed us to identify parameter domains that match the NREM data and, further, enabled description and understanding of both neocortical and hippocampal alternation dynamics with the same model.

We report that neocortical and hippocampal populations are neither endogenously oscillatory nor bistable during NREM sleep, but are *excitable*: each population rests in a stable state from which suprathreshold fluctuations can induce a transient population event that is terminated by the influence of adaptation. Specifically, the neocortex maintains a stable UP state with fluctuation-induced transitions to a transient DOWN state (slow waves), while the hippocampus rests in a stable DOWN state with fluctuation-induced transitions to a transient UP state (SWRs). Under the influence of noise, each region can generate its respective population event spontaneously (due to internally-generated fluctuations) or in response to an external perturbation (such as input from another brain structure). The result is alternations between active and inactive states in both structures with the asymmetric duration distributions observed during NREM sleep (Figure 1D,H). We further observe that variation in the depth of NREM sleep corresponds to variation in the stability of the neocortical UP state. Our findings reveal a unifying picture of the state of hippocampal and neocortical populations during NREM sleep, which suggests that NREM function relies on excitable dynamics in the two regions.

## RESULTS

### UP/DOWN dynamics in an adapting excitatory population model

UP/DOWN alternations are produced in models of neural populations with recurrent excitation and slow adaptive feedback^15-17,20-24^ ^25^. In our model, neuronal population activity is described in terms of the mean firing rate, *r(t)*, subject to a slow negative feedback (i.e. adaptation), *a(t)* (Figure 2A, see Supplemental Info for details).

**Figure 2:**
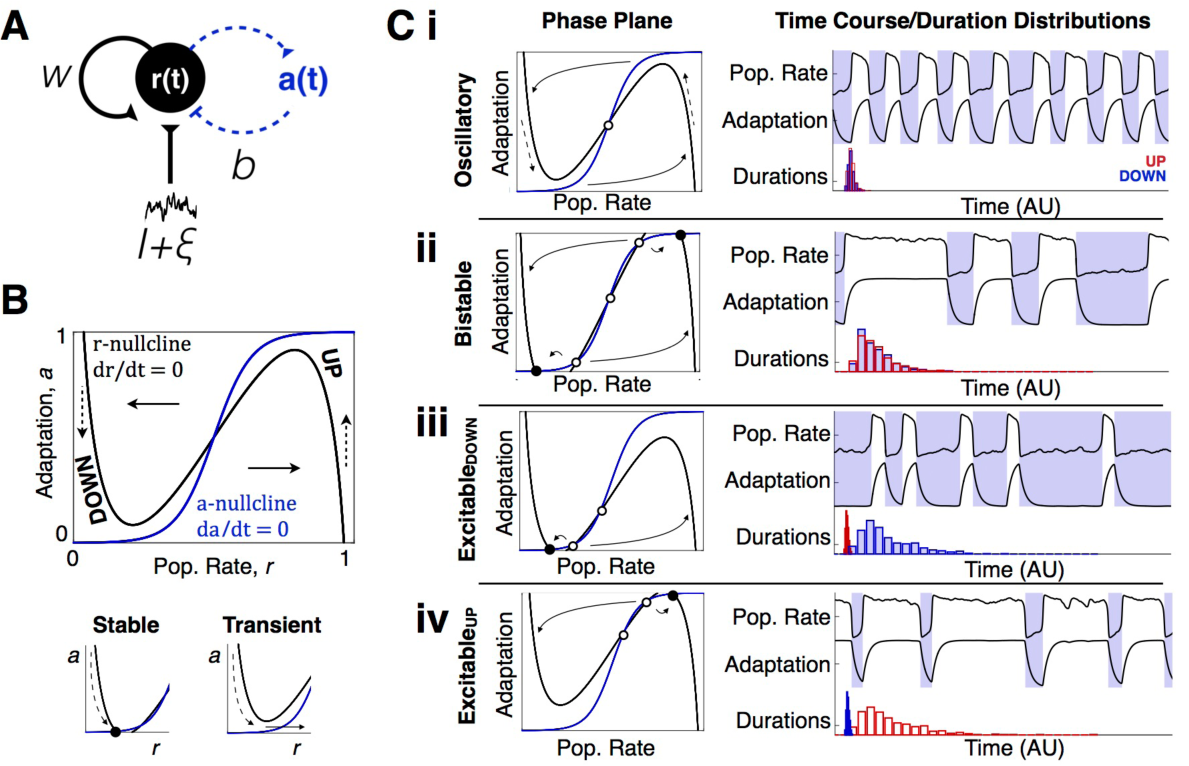
UP/DOWN dynamics in an adapting recurrent neural population. **A:** Wilson-Cowan-like model for a neural population with slow adaptive process. **B:** r-a Phase plane. Model dynamics are seen as trajectories in the phase plane that follow equations 1-2. Dashed arrows indicate slow vertical trajectories at timescale of τ_a_, solid arrows indicate fast horizontal trajectories at timescale of τ_r_. Nullclines (dr/dt = 0, da/dt = 0) and their intersections graphically represent dynamics for a given set of parameter values (parameters as defined in Ci shown). Left and right branches of the r-nullcline correspond to DOWN and UP states, respectively. Stable UP/DOWN states are seen as stable fixed points at nullcline intersections. Transient UP/DOWN states are seen as r-nullcline branch with no intersection. **C:** Four UP/DOWN regimes available to the model, as distinguished by location of stable fixed points (see also Supplemental Figure 3). Representative phase plane (Left), simulated time course and UP/DOWN state duration distributions (right, time units arbitrary) for each regime. Stable fixed points are represented by filled circles, unstable fixed points by empty circles. Parameters: (i-iv) b=1, (i,iii,iv) w=6 (ii) w=6.3, (i) 1=2.5 (ii) 1=2.35 (iii) 1=2.4 (iv) 1=2.6. Default parameters specified in methods.

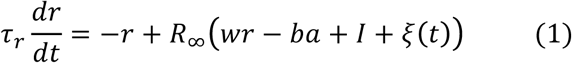

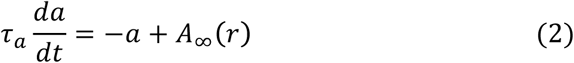

Equations 1-2 describe how *r* and *a* evolve in time as a function of the net input to the population: the sum of the recurrent excitation with weight *w* and a background level of drive with a tonic parameter *I*, and noisy fluctuations *ξ(t)*, minus adaptation weighted by gain parameter *b* (See Supplemental Info for parameter interpretation) *R _∞ (input)_* is the “cellular-level” input-output function, which defines the rate of the population given constant net input. Similarly, *A _∞_ (r)* defines the level of adaptation given a fixed population rate. To enable the analytical treatment of model dynamics in the following section, both *R _∞ (input)_* and *A _∞_ (r)* are taken to be sigmoidal functions.

Model dynamics can be represented as a trajectory in the *r-a* phase plane^26^ (Figure 2B, Supplemental Info), in which steady states, or fixed points, of activity are found at intersections of the *r*-and *a*-nullclines: two curves defined by the conditions 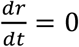 and 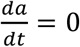 Depending on parameter values, the model can show four distinct regimes of UP/DOWN dynamics - distinguished by whether UP/DOWN transitions are noise- or adaptation-induced, and thus the stability or transient nature of the UP and DOWN states (Figure 2B)^15,16^.

In the oscillatory regime (Figure 2Ci), activity alternates between transient UP and DOWN states at a relatively stable frequency. Adaptation activates during the UP state and brings the population to the DOWN state, during which adaptation inactivates and the population returns to the UP state. Because *r(t)* is fast compared to the slow adaptation, the *r(t)* time course and the phase plane trajectory are square-shaped, with rapid transitions between UP and DOWN states.

If both the UP and the DOWN state are stable, the system is in a bistable regime (Figure 2Cii). In this regime, adaptation is not strong enough to induce UP/DOWN state transitions. However, sufficiently large (suprathreshold) fluctuations can perturb the population activity to cross the middle branch of the *r*-nullcline, resulting in a transition to the opposing branch. Thus, the presence of noise induces alternations between stable UP and DOWN states, resulting in highly variable UP/DOWN state durations.

In the case of a single stable state, the system can still show UP/DOWN alternations in one of two excitable regimes. If the DOWN state is stable (Figure 2Ciii), the system is in an Excitable_DOWN_ regime. The population will remain in the stable DOWN state in the absence of any external influence. However, a brief suprathreshold activating input to the population can trigger a rapid transition to a transient UP state, during which adaptation activates, leading to a return to the DOWN branch. In the presence of noise, UP states are triggered spontaneously by net activating fluctuations. The time course of the model in the Excitable_DOWN_ regime shows long DOWN states of variable durations punctuated by brief stereotyped UP states.

Conversely, if the UP state is stable, the system is in an Excitable_UP_ regime (Figure 2Civ). Brief inactivating input can elicit a switch from the UP state to a transient DOWN state, during which adaptation deactivates, leading to a return to the UP branch. Thus, in the presence of noise, DOWN states are triggered spontaneously by net-inactivating fluctuations. The time course will show longer UP states of variable durations with stereotypically brief DOWN states. These two regimes (Figure 2Ciii,iv) are excitable because relatively small fluctuations in population rate can “excite” the population out of a stable steady state and induce disproportionately large, stereotyped, population events: a transient UP state in the case of the Excitable_DOWN_ regime and a transient DOWN state in the case of the Excitable_UP_ regime, followed by a return to the stable steady state.

### Recurrence, adaptation, and drive control UP/DOWN regimes

How do properties of a neuronal population determine dynamical regime? We use numerical and analytical methods from dynamical systems theory^26^ to reveal how intrinsic and network properties determine the properties of UP/DOWN dynamics in our model. The analysis is summarized here; see Supplemental Info and Supplemental Figures 2-4 for further description of the response properties and parameter dependencies, as well as a discussion of general insights from the mathematical analysis on UP/DOWN dynamics in various physiological contexts.

We first consider how the population rate at steady state, *r*_*ss*_, depends on the level of drive: the *effective* input/output relation (I/O curve) of the recurrently connected population (Figure 3A). If recurrence is weak, the I/O curve increases monotonically with drive and no UP/DOWN alternations are possible. At a critical value of recurrent excitation the population is able to self-maintain an UP state under conditions of reduced drive (Supplemental Figure 2), and UP/DOWN alternations emerge in the I/O curve between a low-rate steady state at weak drive and a high-rate steady state at strong drive. Recurrence and adaptation oppositely influence the dynamical regime at the I/O curve’s center region (Supplemental Figure 3). By solving for parameter values of transitions in the dynamical regime at the half-activation point of the I/O curve (Figure 3B, Supplemental Figure 4), we see that the population will have an oscillatory-centered I/O curve with stronger adaptation (Figure 3B, blue) and a bistable-centered I/O curve with stronger recurrence (Figure 3B, yellow).

**Figure 3:**
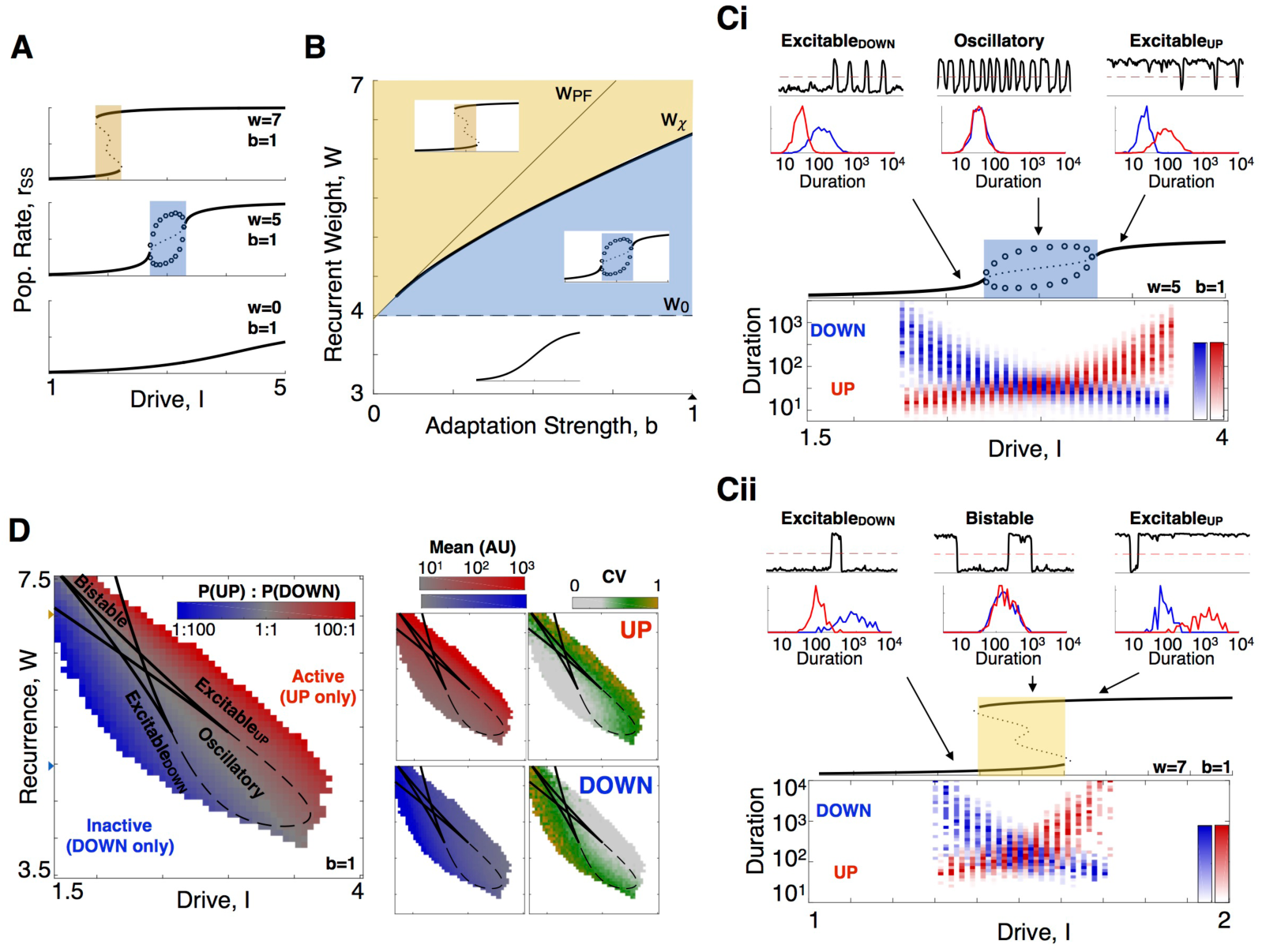
Recurrence, adaptation, and drive determine dynamical regime and duration statistics. **A:** Effective 1/0 curves for populations with increasing levels of recurrent excitation, w. Solid lines indicate stable fixed points, dotted lines indicate unstable fixed points, and circles indicate the upper/ lower bounds of a limit cycle. Yellow indicates bistable regime (2 stable fixed points), blue indicates oscillatory regime (0 stable fixed points with a stable limit cycle). See also Supplemental Figures 3, 4. **B:** Regions in the w-b parameter plane for distinct 1/0 properties, indicated by insets. Blue shading indicates parameter values for a oscillatory-centered 1/0 curve, yellow shading indicates a bistable centered 1/0 curve. Boundary curves correspond to parameters where bifurcations vs I coalesce, see Supplemental Information. W_PF_: pitchfork bifurcation separating 5-fixed point and 3-fixed point (2 stable) regimes. w_x_: degenerate pair of saddle node bifurcations, separating bistable-centered and oscillatory centered 1/0 curves. w_o_: degenerate hopf bifurcation separating oscillatory-centered and monotonic stable 10 curves. **Ci:** Simulated duration distributions on the oscillatory-centered 1/0 curve. (Top) Simulated time course (r vs t) and dwell time distributions (bottom) for low (I=2.6) intermediate (I=3), and high (I=3.4) levels of drive. (Bottom) UP/DOWN state duration distributions as a function of drive to the system. Note logarithmic axis of duration. **Cii:** Same as Ci, for the bistable-centered I/O curve **D:** Duration statistics as a signature of UP/DOWN regime. (Left) Ratio of simulation time in UP and DOWN state. (Right) Mean and CV of simulated dwell times for UP and DOWN states in the l-W parameter space with fixed b (triangle in 3B).

In the absence of noise or external perturbation, only the oscillatory regime will alternate between UP and DOWN states. To illustrate the effects of noise, we consider the effects for the case of an oscillatory-centered I/O curve (Figure 3Ci). Within the oscillatory regime the simulated population rate alternates regularly between transient UP and DOWN states, and UP/DOWN state durations reflect the time scale of adaptation, ∼ τ_a_ (Supplemental Figure 5). For *I*-values above the oscillatory regime, there is a stable UP state fixed point, but noise can evoke transitions to a transient DOWN state (an Excitable_UP_ regime). DOWN state durations still reflect the time scale of adaptation, τ_a_ but UP state durations now reflect the waiting time for random fluctuations to drop the system out of the UP state attractor, and thus vary with noise amplitude (Supplemental Figure 5). For *I-*values further above the oscillatory regime, the effective stability of the UP state increases; DOWN states are less frequent because the larger fluctuations needed to end the UP state are less frequent. Thus, UP states become progressively longer as *I* is increased, while DOWN states stay approximately the same duration (∼ τ_a_). The same case is seen for values of *I* below the oscillatory regime but with UP/DOWN roles reversed (i.e. an Excitable_DOWN_ regime). Similar response properties are seen for the bistable-centered I/O curve (Figure 3Cii). In both cases, the duration distributions plotted vs. drive form a crossed-pair, with a center symmetrical portion (i.e. an oscillatory (Figure 3Ci) or bistable (Figure 3Cii) regime) flanked by the asymmetrical Excitable_DOWN_ and Excitable_UP_ regimes.

In sum, the statistics of UP/DOWN state durations reflect the underlying dynamical regime as seen for simulations in a representative I-w parameter plane (*b*=1, Figure 3D) and in the *I-b* space (*w*=6, Supplemental Figure 5). The mean durations vary continuously over the parameter plane as the level of drive brings the population from a DOWN-dominated to an UP-dominated regime. However, the duration variability (as measured by the coefficient of variation, CV) shows sharp transitions at the boundaries between regimes, which reflect the different mechanism of transitions out of stable and transient states. In general, the durations of stable states are longer and more variable while those of transient states are shorter and less variable, effectively distinguishing oscillatory, bistable, and excitable dynamics.

### Neocortex is in an Excitable_UP_ regime during NREM sleep

The durations of neocortical UP/DOWN states (Figure 1) are indicative of an Excitable_UP_ regime in our model. Neocortical UP states during NREM are longer (1.7±0.92s) compared to DOWN states (0.21±0.05s), and more irregular (CV_UP_ = 1.1±0.27; CV_DOWN_ = 0.38±0.06) (Figure 4A) suggesting a stable UP and transient DOWN state. We directly compared the simulated and experimentally-observed dynamics by matching the statistics of experimental UP/DOWN durations to those in Figure 3D and Supplemental Figure 5. We found that the region of parameter space in which the CV_UP_, CV_DOWN_ and ratio of mean durations is within 2 standard deviations of the experimental durations is in the Excitable_UP_ regime (Figure 4B, Supplemental Figure 7 red outline). We next compared the shapes of the duration distributions between model and experiment. For each model realization (i.e. each point in the *I-w* parameter plane), we calculated the similarity between simulated and experimental duration distributions for each recording session in the experimental dataset (Supplemental Figure 6, 7, Methods). The domain of high similarity between animal data and the model fell in the Excitable_UP_ regime, as indicated by the 25 best fit points and in the average values of similarity (over all 25 sessions) in *I-w* parameter space (Figure 4B) and in the *I-b* parameter space (Supplemental Figure 7). The simulated time course (Figure 4D) and duration distributions (Figure 4C) using the parameter set with highest mean similarity over all sessions revealed a good match between experimental and modeled dynamics. The domain of high similarity was degenerate and remained in the Excitable_UP_ regime with variation in the “fixed” parameters, τ_*a*_, *b*, and the amplitude of the noise (Supplemental Figure 7). We thus found that NREM sleep in the rodent neocortex is characterized by an Excitable_UP_ regime: a stable UP state with noise-induced transitions to a transient DOWN state.

**Figure 4:**
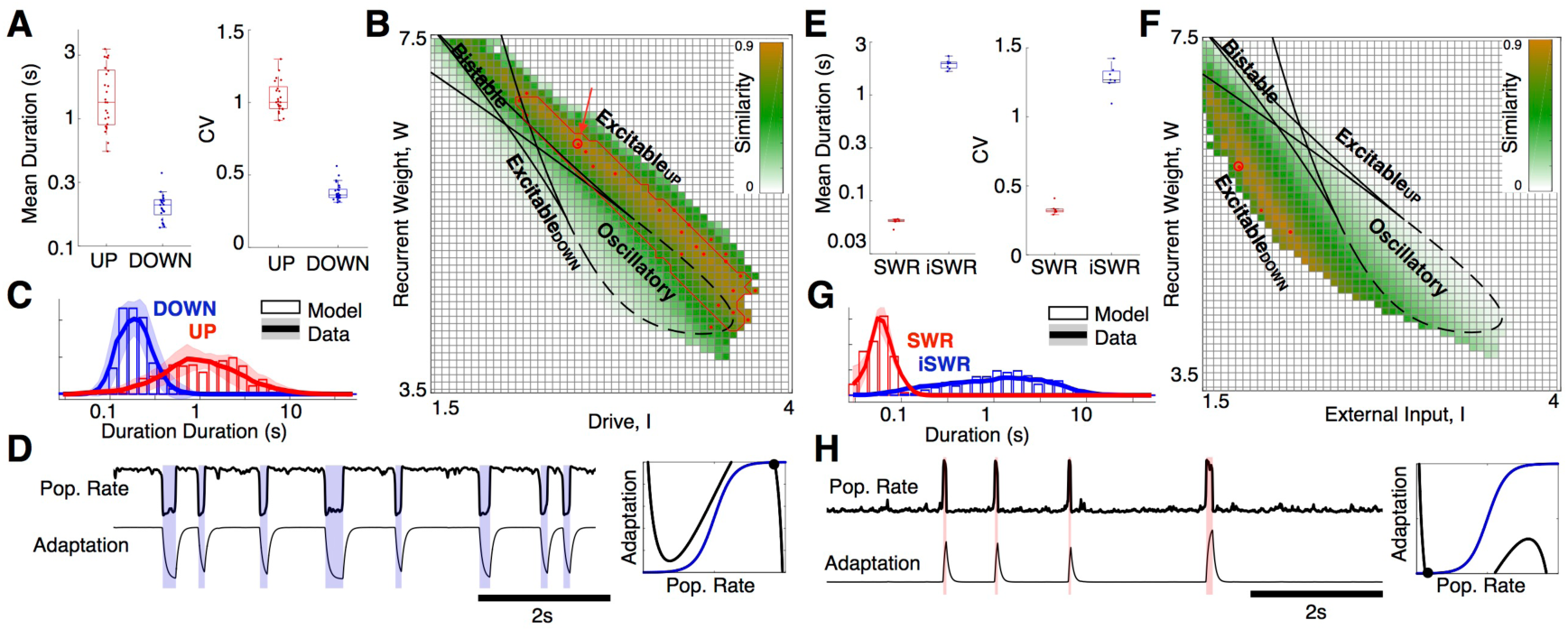
Neocortex and hippocampus are in excitable regimes during NREM sleep. **A:** Mean and CV of UP/DOWN state durations, each point is a single recording session. Center line: media, box limits: upper/lower quartiles, whiskers: data range (excluding outliers). **B:** Similarity between simulated and experimental duration distributions (see Supplemental Figure 6, Methods for similarity calculation). Color indicates mean similarity over all recordings, red line outlines region for which model simulations fall within mean+/-2STD of experimentally observed CV_UP_, CV_DOWN_ and mean_UP_/mean_DOWN_ ratio. 25 red dots indicate most similar parameters for each of the 25 recordings, and large circle (arrow) is best mean similarity. **C:** Neocortical duration distributions for data (mean+/-std over all recordings) and model simulation in the best mean similarity parameters indicated in panel B. (1=2.64, W=6.28, b=1). **D:** Simulated r-a model with the best-matching parameters. (Right) Phase plane diagram for model with best fit parameters, in the Excitable_UP_ regime. **E:** Mean and CV of SWR/interSWR state durations, each point is a single recording session (n=7 hippocampal sessions). **F:** Similarity between simulated and experimental duration distributions as in panel B. G: Hippocampal duration distributions for data (mean +/-std over all recordings) and model simulation in the best mean similarity parameters. **H:** Simulated r-a model with the best-matching parameters indicated in panel F. (1=1.9, W=6, b=1). (Right) Phase plane diagram for model with best fit parameters, in the Excitable _DOWN_ regime.

### Hippocampus is in an Excitable_DOWN_ regime during NREM sleep

Since the burst-like dynamics of SWR is reminiscent of the Excitable_DOWN_ regime of our model, we asked whether these patterns could also be explained by the same principles. InterSWR durations are much longer (mean = 2.0±0.22s) compared to SWR events (mean = 0.06±0.005s), and more variable (CV_InterSWR_ = 1.3±0.10; CV_SWR_ = 0.33±0.04) (Figure 4E) suggesting a stable DOWN and transient UP state (SWR). We applied the duration distribution matching procedure to the SWR/inter-SWR duration distributions and confirmed that the *r-a* model can also mimic SWR dynamics, with a band of high data-model similarity in the Excitable_DOWN_ regime (Figure 4G). Interestingly, our idealized model is not able to capture the short-interval inter-SWR periods associated with occasional SWR “bursts” (Supplemental Figure 7), which suggest the presence of separate SWR-burst promoting mechanisms, possibly arising from interactions with the entorhinal cortex or spatially traveling patterns of SWRs in the hippocampus^27,28^. Accordingly, while the mean ratio and CV_SWR_ of the best fitting model regime were within 2.5 standard deviations of those observed in vivo, the CV of inter-SWR periods was larger than expected from the model (i.e. CV>1). This finding suggests that during NREM sleep the hippocampus is in a stable DOWN-like state, from which internal ‘noise’ or an external perturbation can induce population-wide spiking events.

### Changes in neocortical state correspond to changes in UP state stability

For our initial analysis of the neocortical NREM data we assumed that model parameters were stationary over the course of a sleep session. However, rodent NREM sleep has been classified on a spectrum from light to deep NREM, with higher power in the LFP delta band (1-4Hz) reflecting deeper NREM sleep^29^. To investigate the relationship between changes in cortical state with NREM depth and UP/DOWN dynamics, we calculated the level of delta power in the 8s time window surrounding each UP and DOWN state (Figure 5A). UP state durations varied systematically with delta power (Figure 5A,B,C, Supplemental Figure 8): epochs of lower delta power contained longer UP states, and epochs of higher delta power were associated with shorter UP states (Figure 5A,B,C, Supplemental Figure 8). However, DOWN state durations were invariant with delta power, and the CV of UP state durations was consistently higher than DOWN state durations, as would be expected for Excitable_UP_ dynamics with noise-induced transitions from a stable UP to a transient DOWN state.

**Figure 5:**
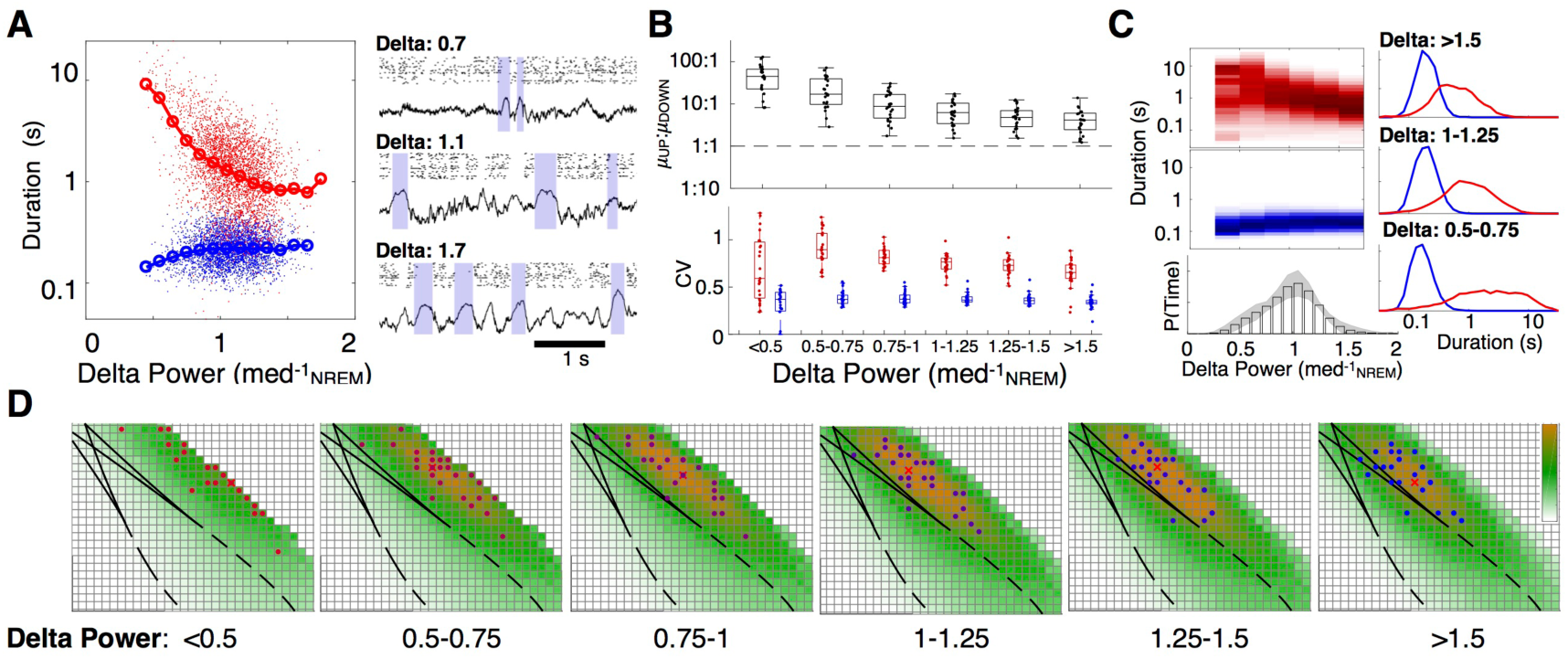
State-dependent variation in alternation dynamics during NREM sleep. **A:** UP/DOWN state durations as a function of delta (1-4Hz) power in a representative recording (delta power normalized to median power during NREM sleep). **B:** Ratio of mean UP/DOWN durations in each neocortical recording as a function of delta power (Top). CV of UP/DOWN durations in each neocortical recording as a function of delta power (Bottom). Center line: media, box limits: upper/lower quartiles, whiskers: data range (excluding outliers). **C:** (Top) Mean distribution of UP and DOWN state durations plotted vs delta power measured in the 8s window around each UP or DOWN state. (Bottom) Mean delta power distribution over all recordings, (error bar: 1 standard deviation). (Right) Mean UP/DOWN state duration distributions for high, medium, and low delta power. **D:** Map of in vivo-model similarity (as in Figure 48, with τ_r_ fixed at τ_r_=5ms), for the delta-power groups as in panel C.

We then grouped the experimental UP/DOWN states by delta power and calculated data-model similarity maps for UP/DOWN state durations in each group (Figure 5D, Supplemental Figure 8). We found that the vast majority of time in all recording sessions was spent in the parameter domain of the Excitable_UP_ regime (Figure 5B, bottom). However, with higher delta power, the best fitting model parameters moved closer to the transition to the oscillatory regime, and the epochs of highest delta power were well-matched by oscillatory dynamics in a small number of sessions.

### Inhibition stabilizes a low-rate UP state and allows perturbation-evoked slow waves

Our previous analyses considered a constant (stationary) source of noise that produced “spontaneous” transitions out of stable states in our model. We now consider a brief input that evokes a transient event. For the hippocampal-like Excitable_DOWN_ regime, a brief *increase* in drive will evoke a transient UP state, (i.e. a SWR, Supplemental Figure 9). In the absence of noise, perturbations must be of sufficient magnitude (i.e. suprathreshold). With noise, the probability to evoke a SWR increases with magnitude of the perturbation (Supplemental Figure 9). A converse situation is apparent for the neocortical-like Excitable_UP_ regime -- a brief *decrease* in drive is able to evoke a transient DOWN state (i.e. a slow wave, Supplemental Figure 9). However, as long-range projections tend to be excitatory, we wondered how an excitatory perturbation might evoke a neocortical UP->DOWN transition.

Neuronal spike rates during the UP state are generally low^18^ with balanced excitatory and inhibitory synaptic inputs^30^. Previous work has shown that models with fast inhibition and slow adaptation can give UP/DOWN alternations in the same four regimes described above^15^ with a low-rate UP state that is stabilized by feedback inhibition^31,32^. We hypothesized that the effect of inhibitory cells may support excitation-induced UP->DOWN transitions, and included an inhibitory population (*τ*_*i*_ *≈ τ*_*e*_) into our model (Figure 6A):

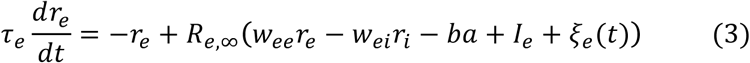

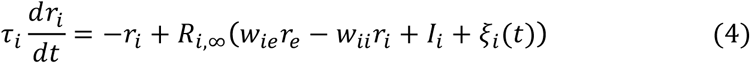

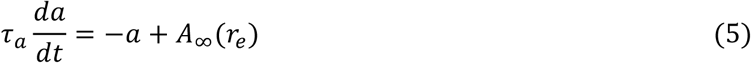

where adaptation acts on the excitatory population and *R*_*e, ∞(input)*_ and *R*_*i, ∞(input)*_ are threshold power law I/O relations, as seen in the in vivo-like fluctuation-driven regime^33^ (Supplemental Info).

**Figure 6:**
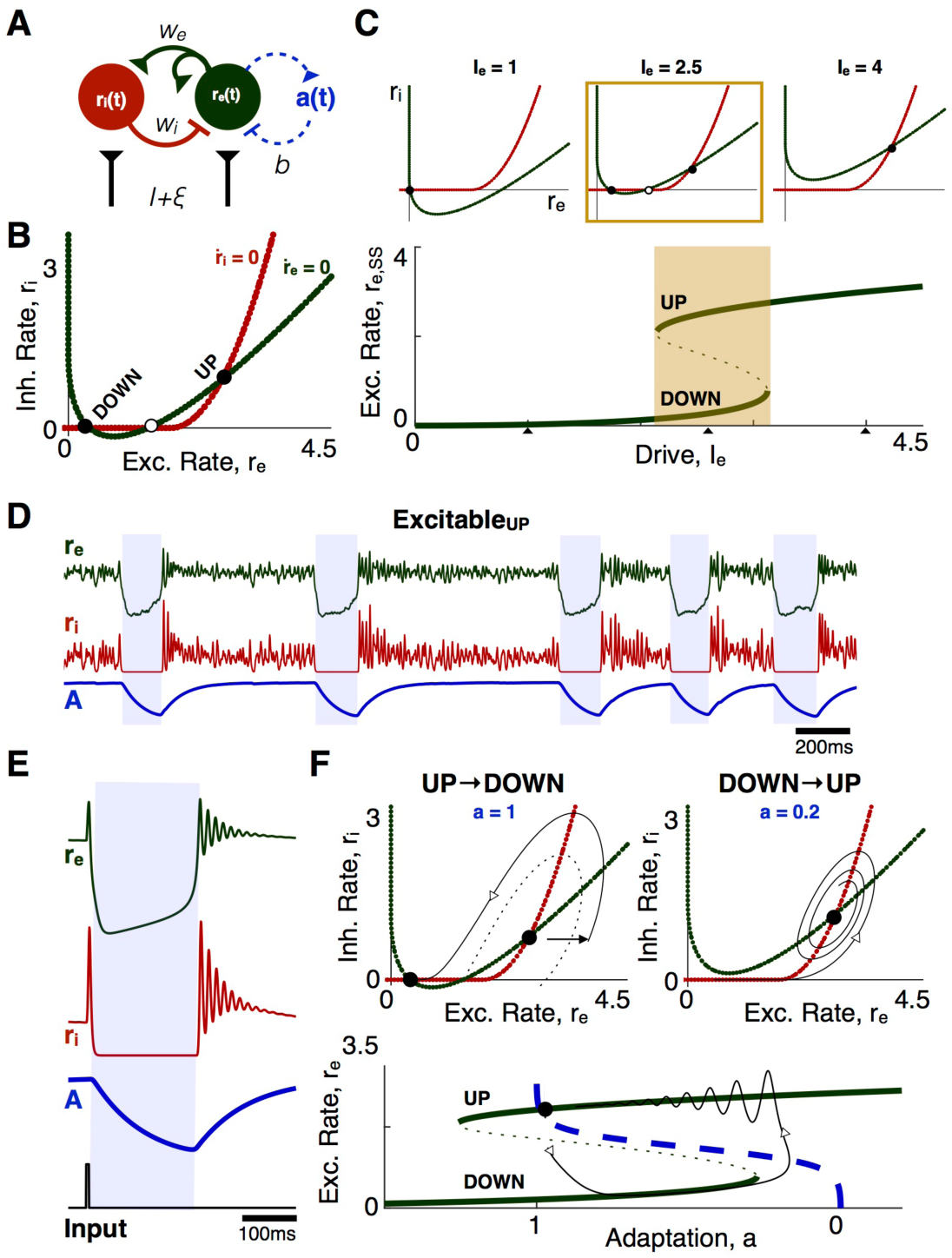
UP/DOWN dynamics in an Adapting Inhibition-Stabilized Network. **A:** Model schematic. **B:** *r*_*e*_-*r*_*i*_ phase plane with *a* frozen. **C:** Bistability in the ISN with *a* frozen. Effective I/O curve of the excitatory population rate, *r*_*e*_, *ss,* with *r*_*e*_-*r*_*i*_ phase planes at low, intermediate, and high levels of drive to the E population (Top). Parameters: *w*_*e*_ = 4, *Wi* = 3. **D:** Excitable_UP_ dynamics in the alSN. Parameters: τ_e_ = τ_i_ *=* 5ms, τ_a_= 200ms **E:** UP/DOWN transitions in the Excitable_UP_ ISN. (Left) Time course of the model in response to brief excitatory input to the excitatory population. **F:** (Bottom) Trajectory of the time course in the *r*_*e*_-*a* phase plane after stimulus-induced transition. The effective I/O curve, r_e, ss_, and the steady state adaptation curve, A_∞_(r), act like the r and a nullclines in the excitation only model. (Top) Trajectories of the stimulus-induced UP->DOWN and subsequent DOWN->UP transition in the *r*_*e*_-*r*_*i*_, phase plane. The dotted line separates the basin of attraction for the DOWN and UP state fixed points.

Given that adaptation is slow we can treat *a* as frozen and visualize model dynamics in the *r*_*e*_*-r*_*i*_ phase plane (Figure 6B). The fixed point value of *r*_*e*_ as a function of drive describes the effective I/O curve of the network (*r*_*ss*_, Figure 6C). Like the excitation-only model, strong recurrent excitation induces bistability at low levels of drive (Supplemental Figure 10). In the bistable condition, the *r*_*e*_*-r*_*i*_ phase plane shows stable UP and DOWN state fixed points, separated by a saddle point (Figure 6B,C). With *a* dynamic, the model can have steady state fixed points on either the UP or the DOWN branch of the I/O curve, resulting in the same regimes as the two-variable model described above^15^ (Supplemental Figure 10).

We investigated Excitable_UP_ dynamics in the adapting, inhibition-stabilized model^33^ (Figure 6D,E,F). Consider a transition from the UP to the DOWN state (Figure 6F). As adaptation slowly de-activates, the system drifts along the DOWN branch. Eventually the DOWN state loses stability, the trajectory reaches and rounds the lower knee of the I/O curve and transitions abruptly to the only remaining stable solution: the UP state. Adaptation then builds as the system returns to the stable UP state fixed point.

Due to the effects of inhibition, small perturbations from the UP state fixed point will exhibit damped, resonant, E-I oscillations as the system returns to the fixed point state. The damped oscillations arise from transient imbalance of excitation and inhibition, and occur when the UP state fixed point is an attracting spiral. As a result, high frequency oscillations (at a time scale set by the excitatory and inhibitory time constants) occur at the DOWN->UP transition. A further implication is that a sufficiently strong *excitatory* input to the excitatory population (Figure 6E) can recruit sufficient inhibition to force the entire network into a DOWN state. This threshold effect is seen as a trajectory in the phase plane that separates the basins of attraction of the UP and DOWN state (i.e. a separatrix, Figure 6F). The separatrix emerges (in reverse time) from the saddle and curves around the UP state fixed point. From this visualization we see that a brief excitatory input to either population can push the trajectory out of the UP state basin of attraction (Figure 6F). Thus, a transient DOWN state (i.e. a slow wave) can be evoked by an excitatory perturbation to either population, as well as due to drops in the excitatory population rate.

## Discussion

To account for cortical dynamics during NREM sleep, we used a firing rate model that represents a neuronal population with positive feedback (recurrent excitation) and slow negative feedback (adaptation). Although the model is idealized, it is amenable to mathematical treatment in terms of a few key parameters and allows us to develop intuitions for the repertoire of dynamics available to an adapting, recurrent neural population. Our analysis of the model revealed how the level of drive and the relative strength of recurrent excitation and adaptation create a spectrum of dynamical regimes with UP/DOWN alternations, defined by the stability or transience of UP and DOWN states (Figure 7A). We found that both neocortical and hippocampal alternations during NREM sleep are well matched by the model in excitable regimes of dynamics that produce characteristically asymmetric distributions of UP and DOWN state durations. We next discuss biological interpretations of the model and implications of the findings for NREM sleep. Additional discussion on general insights of UP/DOWN alternations in other physiological contexts can be found in the Supplemental Info.

**Figure 7:**
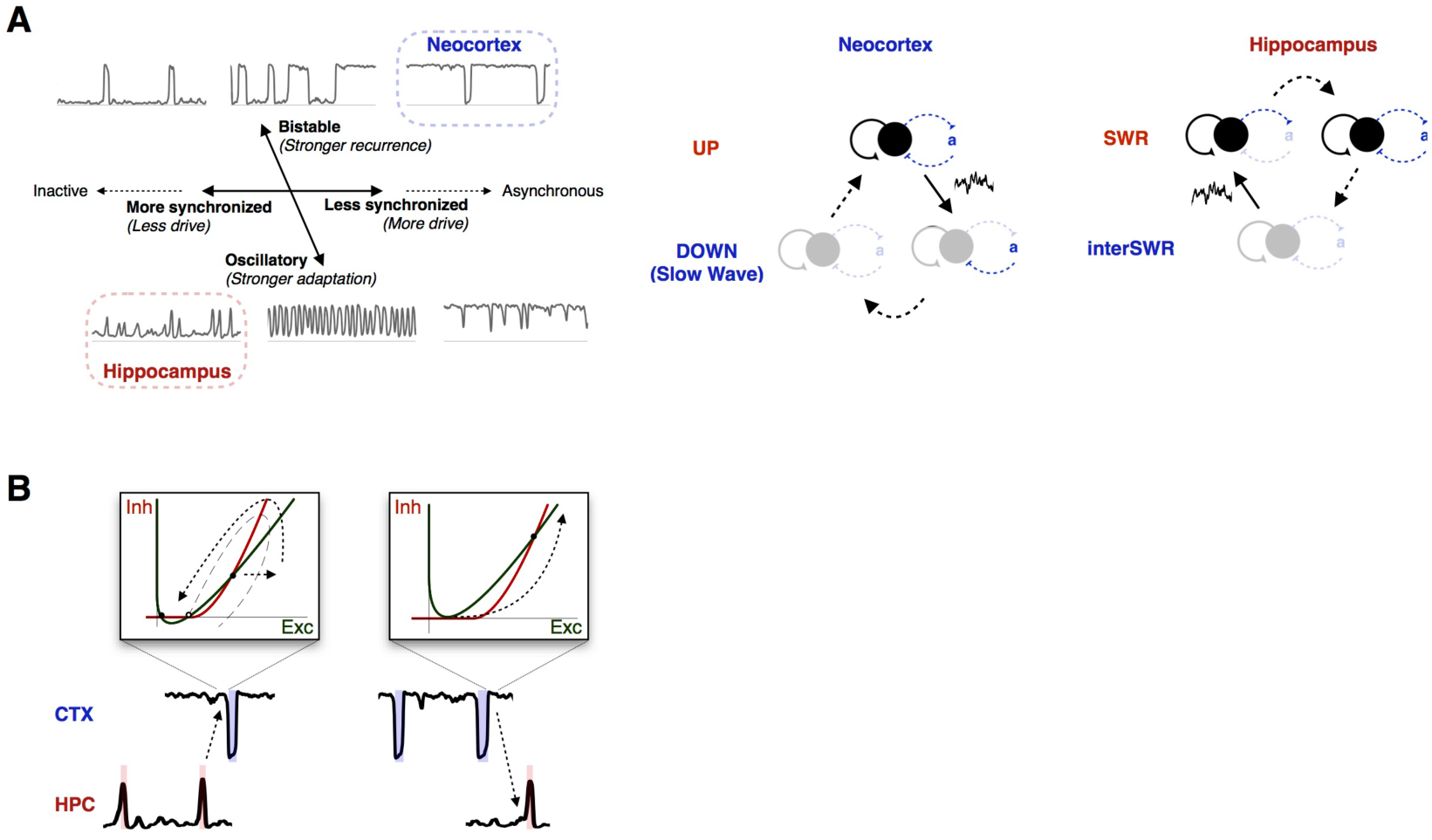
Excitable dynamics of NREM sleep. **A:** UP/ DOWN alternation dynamics are characterized by a spectrum of two effective axes: a degree of synchronization axis determined by the level of drive in the system, and an oscillatory-bistable axis determined by the relative strength of recurrence and adaptation. The tilt of the oscillatory-bistable axis reflects the decrease in the amount of drive needed for equi-duration UP/DOWN states (I _1/2_) as recurrence is increased or adaptation is decreased. (Right) Schematic of the neocortical and hippocampal excitable regimes that produce slow waves and SWRs. Solid arrows indicate noise-induced transitions. **B:** Hypothesis for mechanism of SWR/slow wave coupling by excitable dynamics.

### UP/DOWN dynamics of the neocortical NREM slow oscillation

Despite the widely used term slow “oscillation”^1^, the asymmetric duration distributions during NREM predict that the NREM slow oscillation is reflective of Excitable_UP_ dynamics: an aperiodic process in which activity fluctuations during stable UP states can lead to transient DOWN states (slow waves). A key feature of the model is the noise responsible for initiating spontaneous UP->DOWN transitions. In neuronal network modeling, “noise” often refers to unidentified fluctuations in physiological activity. Broadly, biological noise can be divided into fluctuations internal to the population and fluctuations from afferent projections. While we do not explicitly distinguish them in the model, we assume that both sources play a role in initiating cortical UP->DOWN transitions. Population rate fluctuates during the UP state due to finite size effects^34^ and temporal correlations that emerge with strong recurrent connections^35^. Similarly, the level of afferent activity from thalmo- or cortico-cortical projections would be expected to fluctuate. We also note that while the isolated cortex can produce UP/DOWN state alternations^36^, we should consider the thalamocortical system for an understanding of slow wave dynamics in vivo^37^. Because the cortex and corresponding thalamic nuclei are highly interconnected, cortex and thalamus may transition UP and DOWN together and reflect interacting (as opposed to independent) systems. However, it was recently found that cortex tends to lead the thalamus into the DOWN state^38^. Future work should expand the model to include a thalamic population, which would also allow a better understanding of the interaction of slow waves with thalamocortical spindle oscillations^8,39,40^.

We found that the depth of NREM sleep reflects an evolution of the stability of the UP state in a manner that resembles the stages of NREM/SWS sleep in humans^41^. In light NREM sleep (N1 stage, using the human clinical term), long UP states are occasionally punctuated by neuronal silence-associated positive delta or slow waves, which can be localized at one or few recording sites across the cortical mantle^8^. As sleep deepens, the incidence of DOWN states increase and they become synchronous over larger cortical areas^42^ (N2 stage). The DOWN-UP transitions occasionally become strongly synchronous, producing a sharp LFP wave, known as the K complex^43^. With further deepening of sleep, DOWN states become more frequent and short episodes of repeating DOWN states may become rhythmic (N3 stage). While direct comparison between rodent and human sleep data was not performed, we found a similar evolution in rodent NREM. Quantifying the time spent in these sub-states revealed that the N3-like oscillatory state in the rat occupies only a small fraction of NREM sleep, whereas in humans this stage is more prominent. Our model predicts that the stages of sleep reflect different stability of the UP state, which may be due to 1) decreased recurrent strength, 2) decreased neuronal excitability or 3) increased strength of adaptation^44^.

While the model is ambiguous to the biophysical substrate of adaptation, we can make some predictions: first, the adaptive process responsible for neocortical UP/DOWN alternations should be constitutively active during the UP state and deactivate during the hyperpolarized DOWN state. Subthreshold adaptation conductances are a feasible candidate, given that most neurons are silent or fire at a very low rate during any given UP state. Adaptation in our model could also include effects of hyperpolarization-activated excitatory processes, such as the h-current. Second, the neocortical adaptive process should recover at a time scale reflective of the DOWN state duration (∼200ms). Our model can be used to predict how changing properties of the relevant adaptive mechanisms experimentally or in disease should change the duration statistics of the slow oscillation, which can guide for future experiments to uncover the biophysical substrates of the neocortical slow oscillation.

### On differential neocortical and hippocampal dynamics

Neocortical slow oscillations and hippocampal SWRs are present simultaneously during NREM sleep. Although they appear fundamentally different, our model with just a few parameters can quantitatively account for both by using different parameter values. In the hippocampus, the inter-SWR period is not entirely inactive, but maintains a low rate of activity. SWR-initiating noise could be fluctuations in ongoing low-rate activity during the iSWR period or fluctuations in drive from the entorhinal cortex. The different durations of the transient neocortical DOWN state and hippocampal SWR indicate that alternation dynamics in the two regions are most likely mediated by different adaptive processes, and that the hippocampal adaptive process should activate on a time scale that reflects the SWR durations (∼60ms). Previous work has revealed threshold behavior in the generation of SWRs, indicative of Excitable_DOWN_ dynamics, with a GABA_B_-mediated adaptation mechanism^46,47^. Our model suggests that a stronger adaptive process in the hippocampus would favor Excitable_DOWN_ dynamics.

As the relevant parameter is the *relative* strength of adaptation and recurrence, the different nature of recurrent connectivity in the two regions may also be responsible for their differing dynamics. Strongly recurrent pyramidal cell populations are found in neocortical layer 5 and the hippocampal CA2 and CA3a subregions ^48^, the loci of UP state and sharp wave initiation, respectively^49,50^. However, crucial differences exist between connectivity of neocortical layer 5 and hippocampal CA2-3 regions. The neocortex is a modularly organized structure. In contrast, the hippocampus can be conceived as a single expanded cortical module^51^. Excitatory connectivity in layer 5 is local (200 *µ*m), dense (up to 80% connection probability), and follows a ‘Mexican hat’ excitatory-inhibitory spatial structure with strong local excitatory connections and spatially extensive inhibition^52^. In contrast, excitatory connectivity in the hippocampus is sparse and spatially extensive^48^, with local inhibitory connections^53,54^. While layer 5 excitatory synapses are relatively strong, the transmitter release probability of synapses between hippocampal pyramidal neurons is very low, resulting in comparatively weak synapses^55^. Together, these factors indicate that the effective strength of recurrence in the hippocampus is lower than that in neocortex, which would result in DOWN-dominated as opposed to UP-dominated dynamics, as are observed. To further understand the physiological factors responsible for the distinct NREM dynamics in the two regions will require experimental manipulations that independently manipulate adaptation, recurrent excitation, and excitability.

### NREM function through stochastic coordination of excitable dynamics

According to the two-stage model of memory consolidation^56,57^, the hippocampus acts as a fast, but unstable, learning system. In contrast, the neocortex acts as a slow learning system that forms long-lasting memories after many presentations of a stimulus. The two-stage model proposes that recently-learned patterns of activity are reactivated in the hippocampus during SWRs, which act as a “training” signal for the neocortex, and that the neocortical consolidation of those patterns relies on SWR-slow wave coupling^5,58^. Excitable dynamics provide a mechanism for coordination of slow waves and SWRs (Figure 7B): the excitatory kick of a hippocampal SWR can induce a neocortical UP->DOWN transition by briefly disrupting the neocortical excitatory/inhibitory balance, while the population burst at the neocortical DOWN->UP transition can induce a hippocampal SWR.

Extensive experimental evidence points towards temporal coordination between slow waves and SWRs. Slow waves in higher-order neocortical regions are more likely following SWRs^5,6^, and SWR->slow wave coupling is associated with reactivation in the neocortex^5,58,59^. As is observed in vivo, the ability of a transient input to evoke a slow wave in our model is probabilistic, due to an interaction between the magnitude of perturbation, local noise, and the stability of the UP state. The probability of SWR->slow wave induction likely varies by brain state, cortical region, and even SWR spiking content. Further work to investigate how these factors shape SWR->slow wave coupling will likely shed light on the brain-wide mechanisms of memory consolidation.

How then, does a SWR-induced neocortical slow wave induce changes in the neocortex? Recent work has found that SWR->slow wave coupling alters spiking dynamics at the subsequent neocortical DOWN->UP transition^58^ (aka the k complex), which acts a window of opportunity for synaptic plasticity that supports NREM functions^10,60-62^. Interestingly, the interaction between excitation and inhibition produces a transient (gamma-like) oscillation at the DOWN->UP transition in our model. This brief oscillation is reminiscent of the gamma (∼60-150Hz) activity following slow waves *in vivo*^18^ and may act to coordinate and promote plasticity between cell assemblies^63^.

In turn, the burst of neocortical activity during the k complex could induce a SWR in the hippocampus. The functional role of slow wave->SWR coupling is less well understood, but hippocampal SWRs are more likely immediately following slow waves in some neocortical regions - including the entorhinal cortex^5,7^. Slow wave->SWR coupling could provide a mechanism by which neocortical activity is able to bias SWR content, or another mechanism by which the SWR could bias neocortical activity at the DOWN->UP transition. Further, a SWR-slow wave-SWR loop could produce the occasional SWR bursts not captured by our model of hippocampal SWR activity in isolation. Future work on regional or state-dependent differences in the directionality of slow wave-SWR coupling could provide insight into the physiological mechanisms that support memory consolidation.

## Conclusions

Our results reveal that NREM sleep is characterized by structure-specific excitable dynamics in the mammalian forebrain. We found that a model of an adapting recurrent neural population is sufficient to capture a variety of UP/DOWN alternation dynamics comparable to those observed *in vivo*. The neocortical “slow oscillation” is well-matched by the model in an Excitable_UP_ regime in which a stable UP state is punctuated by transient DOWN states, while the hippocampal sharp waves are well-matched by the model in an Excitable_DOWN_ regime in which a stable DOWN state is punctuated by transient UP states (Figure 7A). These complementary regimes of excitable dynamics allow each region to produce characteristic slow wave/SWR events spontaneously or in response to external perturbation. Our results offer a unifying picture of hippocampal and neocortical dynamics during NREM sleep, and suggest a mechanism for hippocampal-neocortical communication during NREM sleep.

## Supporting information

## Acknowledgements

The authors would like to thank Rachel Swanson, William Muñoz, Brendon Watson, Andres Grosmark for discussions during the development of the project and extensive feedback on the manuscript, the NIH training grant for computational neuroscience T90DA043219 for funding and the TPCN trainees for their feedback on the manuscript, and Brendon Watson and Andres Grosmark for generously sharing their data.

## Author Contributions

Conceptualization, DL GB JR; Methodology, DL JR; Formal Analysis, DL JR; Writing – Original Draft, DL JR; Writing – Review & Editing, DL, GB, JR; Supervision, GB and JR.

## METHODS

### Datasets

The datasets used were reported in Watson et al 2016 (neocortex) and Grosmark and Buzsaki 2016 (hippocampus), and are briefly summarized here.

For the cortical dataset, silicon probes were implanted in frontal cortical areas of 11 male Long Evans rats. Recording sites included medial prefrontal cortex, anterior cingulate cortex, premotor cortex/M2, and orbitofrontal cortex. Neural activity during natural sleep-wake behavior was recorded using high-density silicon probes during light hours in the animals’ home cage. 25 recordings of mean duration 4.8+/-2.2hrs were recorded. The raw 20kHz data was low-pass filtered and resampled at 1250Hz to extract local field potential information. To extract spike times, the raw data high-pass filtering at 800Hz, and then threshold-crossings were detected. KlustaKwik software was used to cluster spike waveforms occurring simultaneously on nearby recording sites, and Klusters software was used for manual inspection of waveforms consistent with a single neuronal source. Units were classified into putative excitatory (pE) and putative inhibitory (pI) based on the spike waveform metrics. Each animal had 35+/-12 detected pE units and 5+/-3 detected pI units on average.

For the hippocampal dataset, silicon probes were implanted in the dorsal hippocampus of 4 male Long Evans rats (7 recordings total). Neural activity during sleep was recorded before and after behavior on a linear track. LFP and spikes were extracted similar to the cortical dataset. Sharp-wave ripple events were detected as described in Grosmark and Buzsaki 2016, with 3134-11898 SWRs detected per recording and used for subsequent analysis.

### NREM Detection

Sleep state was detected using an automated scoring algorithm as described previously (Watson et al 2016), with some modifications. As only the NREM state was used in this study, we describe here the process for NREM detection. However, the code for full state detection is available at https://github.com/buzsakilab/buzcode. NREM sleep was detected using the FFT spectrogram of a neocortical LFP channel, calculated in overlapping 10s windows at 1s intervals. Power in each time window was calculated for frequencies that were logarithmically spaced from 1 to 100Hz. The spectral power was then log transformed, and z-scored over time for each frequency. The slow wave power (signature of NREM sleep) was calculated by weighting each frequency by a weight determined from the mean of the weights for the first principal components from the dataset in Watson et al 2016, which was found to distinguish NREM and non-NREM in all recordings. While the same dataset was used here, using the filter (i.e. weighted frequency)-based approach as opposed to PCA makes the algorithm robust for a wider range of recording conditions, especially those in which there is less time spent asleep (and thus NREM may not be expected to account for the largest portion of variance). Like the first principal component, the slow wave filtered signal was found to be bimodal in all recordings, and the lowest point between modes of the distribution was used to divide NREM and non-NREM epochs.

In the hippocampal dataset, manual NREM scoring as reported in Grosmark and Buzsaki 2016 was used for this study.

### Slow Wave Detection

Slow waves were detected using the coincidence of a two-stage threshold crossing in two signals (Supplemental Figure 1A,B): a drop in high gamma power (100-400Hz, representative of spiking (Watson et al 2017)) and a peak in the delta-band filtered signal (0.5-8Hz). The gamma power signal was smoothed using a sliding 80ms window, and locally normalized using a modified (non-parametric) Z-score in the surrounding 20s window, to account for non-stationaries in the data (for example due to changes in brain state and noise), that could result in local fluctuations in gamma power. The channel used for detection was determined as the channel for which delta was most negatively correlated with spiking activity, while gamma was most positively correlated with spiking activity.

Two thresholds were used for event detection in each LFP-derived signal, a “peak threshold” and a “window threshold”. Time epochs in which the delta-filtered signal crossed the peak threshold were taken as putative slow wave events, with start and end times at the nearest crossing of the window threshold. Peak/window thresholds were determined for each recording individually to best give separation between spiking (UP states) and non-spiking (DOWN states) (Supplemental Figure 1C). To determine the delta thresholds, all peaks in the delta-filtered signal greater than 0.25 standard deviations were detected as candidate delta peaks and binned by peak magnitude. The peri-event time histogram (PETH) for spikes from all cells was calculated around delta peaks in each magnitude bin, and normalized by the mean rate in all bins. The smallest magnitude bin at which spiking (i.e. the PETH at time = 0) was lower than a set rate threshold (the “sensitivity” parameter, Supplemental Figure 1D) was taken to be the peak threshold. For example, a sensitivity of 0.5 means that the delta peak threshold is set to the smallest threshold for which spiking drops below 50% of mean spiking activity. The window threshold was set to the average delta value at which the rate crosses this threshold in all peak magnitude bins. The gamma thresholds were calculated similarly, but using drops below a gamma power magnitude instead of peaks above a delta magnitude.

Once the thresholds were calculated, candidate events were then detected in the delta and gamma power signals, and further limited to a minimum duration of 40ms. Slow wave events were then taken to be overlapping intervals of both the gamma and delta events. DOWN states with spiking above the sensitivity threshold were thrown out.

Detection quality was checked using a random sampling and visual inspection protocol. LFP and spike rasters for random 10s windows of NREM sleep were presented to a manual scorer, who marked correct SW detections, false alarms, and missed SWs. This protocol was used to estimate the detection quality (miss %, FA %) for each recording (Supplemental Figure 1E), and to optimize the detection algorithm. 1,085 - 21,147 slow waves (i.e. UP/DOWN states) were detected per recording and used for subsequent analysis.

### Model Implementation

Phase plane and bifurcation analysis of the model in the absence of noise was implemented in XPP, and a similar code was implemented in MATLAB for simulations of the model with noisy input, for the analysis of UP/DOWN state durations. Noise was implemented using Ornstein-Uhlenbeck noise.

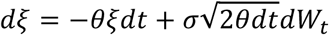

where *W*_*t*_ is a Weiner process. Time scale *θ* = and standard deviation *σ*= were used unless otherwise specified.

Simulations of equations [1-2] and [3-5] were performed in Matlab using the ode45 solver, with input noise *ξ(t)* pre-computed independently for each simulation using forward Euler method with time step dt=0.1. Accuracy was assessed by comparing results for time steps dt=0.1 and dt=0.05 for a subset of simulations. Statistics for simulations with noise were determined by simulations of duration 60,000 (AU).

A simulated time course was determined to have UP/DOWN states if the distribution of r(t) was bimodal, as determined using a hartigans dip test (Hartigan and Hartigan 1985, implementation at http://www.nicprice.net/diptest/). UP/DOWN state transitions were detected as threshold crossings between high and low rate states. To avoid spurious transition detection due to noise, a “sticky” threshold was used: the threshold for DOWN->UP transitions was taken to be the midpoint between positive crossings of a threshold between the high rate peak of the rate distribution and the inter-peak trough, while the threshold for UP->DOWN transitions was the midpoint between the low rate peak of the rate distribution and the inter-peak trough.

All simulation and analysis code is available at https://github.com/dlevenstein/Levensteinetal2018.

### UP/DOWN State Duration Matching

In vivo and simulated UP/DOWN state durations were compared using a non-parametric distribution matching procedure (Supplemental Figure 6). Similarity was calculated as

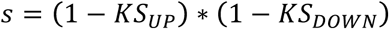

where

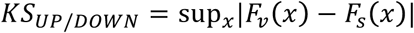

is the Kolmogorov-Smirnov (KS) statistic, in which sup_x_ is the supremum function and *F*_*s/v*_*(x)* are the empirical cumulative distributions of simulated and in vivo durations. In short, KS measures the largest difference between the observed cumulative distributions for simulated and in vivo durations, where *KS*_*UP*_ = 0 indicates that the in vivo/simulated UP state durations distributions are identical and *KS*_*UP*_ = 1 indicates that the in vivo/simulated DOWN state durations distributions are non-overlapping. Similarity is thus bounded between 0 and 1, where *s =* 1 indicates that both UP and DOWN state distributions are identical between simulation and the experimental observation, and *s =* 0 indicates that either the observed UP or DOWN state distributions are non-overlapping with the modeled durations.

There is one free parameter in the fitting procedure, which is *τ* the population time constant, or equivalently, the time scale factor from non-dimensionalized model time and seconds. For each simulation, we tested time scale factors from 1ms to 25ms with increments of 0.1ms and used the time scale parameter that gave the highest value for *s*, thus preserving the shapes of the distributions and the relative values of UP/DOWN state durations.

