## Supplementary material for "Excitable dynamics of NREM sleep: a unifying model for neocortex and hippocampus"

CONTENTS

1. *Description of the r-a model*
2. *Description of the model dynamics in the r-a phase plane*
3. *UP/DOWN bistability in the adaptation-free population*
4. *Bifurcation analysis of the model*
5. *Physiological Interpretation of Model Parameters*
6. *General Insight to UP/DOWN dynamics*
7. *Description of the e-i-a model*

*Description of the Model*

The r-a model represents the mean firing rate or activity of a neural population with activity-driven adaptation a(t).

$$\tau_r \dot{r} = -r + R_\infty\big(wr - ba + I + \xi(t)\big) \qquad (1)$$

$$\tau_a \dot{a} = -a + A_\infty(r) \qquad (2)$$

with activation functions

$$R_\infty(x) = \frac{1}{1 + e^{-(x - x_0)}}$$

and

$$A_\infty(r) = \frac{1}{1 + e^{-k(r - r_0)}}$$

Unless otherwise specified, we use $x_0 = 5$, $r_0 = 0.5 = R_\infty(x_0)$, and $k = 15$ to parameterize the activation functions. Values of other parameters are as indicated in the text and figure legends. The rate and adaptation variables can be considered as non-dimensionalized, scaled by their maximum possible values; similarly, we use $\tau_r = 1$ (unless otherwise specified) so that time is dimensionless (AU, arbitrary units), scaled by

[1] Center for Neural Science, New York University
[2] New York University Neuroscience Institute
[3] Courant Institute for Mathematical Sciences, New York University

the time constant for firing rate.

Under constant net input, the population rate will approach a steady state level given by the input-output relation, $R_\infty(input)$. Adaptation is similarly activated by neuronal activity to a steady state $A_\infty(r)$. For mathematical convenience, $R_\infty(input)$ and $A_\infty(r)$ are taken to be sigmoidal functions. The time constants $\tau_r$ and $\tau_a$ determine how quickly $r(t)$ and $a(t)$ will approach their steady state values and time has been non-dimensionalized to arbitrary model units (AU) so $\tau_r = 1$.

*Description of model dynamics in the r-a phase plane*

Model dynamics can be represented as a trajectory in the $r$-$a$ phase plane (Figure 2B, [1]). In the phase plane, trajectories are predictable from Steady states, or fixed points, of activity are found at intersections of the nullclines and may be either stable (attractors) or unstable. The $r$-nullcline is N-shaped with a right branch at $r{\sim}1$ and a left branch at $r{\sim}0$, which correspond to UP and DOWN states of activity (Figure 2B). When adaptation is slow (i.e. $\tau_a \gg \tau_r$), trajectories move horizontally toward the UP/DOWN branches with time scale $\tau_r$. At the UP/DOWN branches, trajectories drift along the $r$-nullcline as adaptation activates or inactivates with the time scale $\tau_a$. If the branch contains a stable fixed point, the system will remain in the UP or DOWN state until some perturbation induces a transition to the opposing branch. If there is no fixed point, the trajectory transitions to the opposing branch at the turning point of the $r$-nullcline (Figure 2B). In this way, an UP or DOWN state in the model can be either stable: requiring a perturbation to evoke a transition to the opposite state, or transient: automatically transitioning to the opposite state due to the activation or inactivation of adaptation.

*UP/DOWN bistability in the adaptation-free population*

UP/DOWN alternations are possible only if the population can potentially exist in an UP or a DOWN state. This requires adequate strength of recurrent excitation, $w$, to self-maintain the UP state under conditions of low drive. We show this first for a reduced case without adaptation dynamics ($b = 0$). In this case, the population rate $r(t)$ satisfies

$$\frac{dr}{dt} = -r + R_\infty(wr + I)$$

The phase space is reduced from a plane to a line, and dynamics of the population rate correspond to motion along the $r$-axis following eqn. 3 (Supplemental Figure 2A). The motion is rightward (rate increasing) where $dr/dt > 0$ and leftward where $dr/dt < 0$. If recurrent excitation is sufficiently strong, the graph of $dr/dt$ vs. $r$ is N-shaped and there can be two stable (and one unstable) fixed points: an UP state of high activity at $r \approx 1$ and a DOWN state of low activity at $r \approx 0$.

The population rate at fixed points, $r_{ss}$, depends on the level of drive, $I$, as is described by the *effective* input/output relation (I/O curve) of the recurrently connected population (Supplemental Figure 2B). If recurrence is weak ($w = 0$), the I/O curve increases monotonically with $I$. With increased recurrence, the I/O curve shows a central region of bistability between a low-rate fixed point at weak drive and a high-rate fixed point at strong drive. In the $I$-$w$ parameter space (Supplemental Figure 2C), the bistable region (yellow) has borders that correspond to saddle-node bifurcations at the knees of the I/O curve. UP/DOWN bistability emerges at a critical value of recurrence ($w = 4$), at a level of drive for which the unconnected population would be minimally activated ($I = I_0 - 2$, where $R_\infty(I_0) = 0.5$, see Methods). Consequently, a first general insight of recurrent population rate models is that UP/DOWN bistability will emerge in neuronal populations with sufficiently strong recurrent excitation, during conditions of low drive.

*Bifurcation analysis of the model*

With adaptation dynamics reintroduced ($b = 1$), the mechanism for UP/DOWN bistability is like that in the adaptation-free case. Increasing recurrence (larger $w$) enhances the N-shape of the $r$-nullcline, allowing for multiple stable fixed points at intersections of the $r$- and $a$-nullclines (Supplemental Figure 3A). The effective I/O curve is again bistable-centered for strong recurrence (Figure 3A, top). The region of $I$-$w$ parameter space with multiple fixed points is now "butterfly-shaped" (Supplemental Figure 3D), with a bistable regime again inside the yellow region at higher values of $w$.

In contrast to the effect of $w$, stronger adaption (larger $b$) diminishes the N-shape of the $r$-nullcline and decreases likelihood for multiple fixed points (Supplemental Figure 3B). As a result, the network can now oscillate at intermediate values of $w$, for which the I/O curve appears oscillatory-centered (Figure 3A, middle). The oscillatory region in $I$-$w$ parameter space is blue, with borders corresponding to Hopf bifurcations (Supplemental

Figure 3D). Increasing the strength of adaptation increases the domain of this oscillatory region in $I$-$w$ parameter space (Supplemental Figure 3F).

Increasing drive raises the $r$-nullcline and brings the population from a stable DOWN state at low drive to a stable UP state at high drive, with fixed points that trace out the I/O curve (Supplemental Figure 3C). The effective I/O relations, accounting for noise, are represented by statistical properties of UP/DOWN state durations (Figure 3C). The duration distributions plotted vs. drive form a crossed-pair, with a center symmetrical portion (i.e. an oscillatory (Figure 3Ci) or bistable (Figure 3Cii) regime) flanked by the asymmetrical $\text{Excitable}_{\text{DOWN}}$ and $\text{Excitable}_{\text{UP}}$ regimes. A complementary view of the I/O properties is the graph of decreasing fraction of time in the DOWN state as drive is increased. This Prob(DOWN), or silence density, has been used as an experimental metric of the degree of cortical synchronization [2]. By this terminology, more synchronized regimes correspond to more time spent in the DOWN state, and less synchronized regimes correspond to more time spent in the UP state. Increased drive eventually leads to an UP-only (asynchronous) regime. Thus, the level of drive defines a spectrum from more synchronized to less synchronized dynamics in the model defines a set of characteristic axes in the "space" of UP/DOWN alternation dynamics.
We next describe an analysis of the dynamic regime at the I/O curve's center region, which reveals how the relative strength of recurrence and adaptation form a spectrum from bistable to oscillatory dynamics (Figure 3B).

To analyze the parameter space of the model, we used standard procedures from dynamical systems theory (Strogatz). To determine the linear stability and class of a fixed point ($\dot{r} = \dot{a} = 0$) at $[r^*,\ a^*]$, we evaluate the eigenvalues of the Jacobian matrix

$$J = \begin{bmatrix} \frac{\partial \dot{r}}{\partial r} & \frac{\partial \dot{r}}{\partial a} \\ \frac{\partial \dot{a}}{\partial r} & \frac{\partial \dot{a}}{\partial a} \end{bmatrix}_{[r^*\ a^*]} = \begin{bmatrix} -1 + \frac{\partial R_\infty(X)}{\partial r} & \frac{\partial R_\infty(X)}{\partial a} \\ \frac{1}{\tau}\frac{\partial A_\infty(r)}{\partial r} & -\frac{1}{\tau} \end{bmatrix}_{[r^*\ a^*]}$$

where $X = wr - ba + I$ is the total input to the population. Simplifying the partial derivatives as

$$\frac{\partial R_\infty(X)}{\partial r} = \frac{\partial R_\infty(X)}{\partial X}\frac{\partial X}{\partial r} = w\frac{\partial R_\infty(X)}{\partial X}$$

$$\frac{\partial R_\infty(X)}{\partial a} = \frac{\partial R_\infty(X)}{\partial X}\frac{\partial X}{\partial a} = -b\frac{\partial R_\infty(X)}{\partial X}$$

gives the Jacobian for an arbitrary fixed point:

$$J = \begin{bmatrix} -1 + w\dfrac{\partial R_\infty(X)}{\partial X} & -b\dfrac{\partial R_\infty(X)}{\partial X} \\ \dfrac{1}{\tau}\dfrac{\partial A_\infty(r)}{\partial r} & -\dfrac{1}{\tau} \end{bmatrix}_{[r^* \; a^*]}$$

where

$$\frac{\partial R_\infty(X)}{\partial X} = \frac{e^{-(X-x0)}}{(e^{-(X-x0)}+1)^2} = R_\infty(X)(1 - R_\infty(X))$$

$$\frac{\partial A_\infty(r)}{\partial r} = \frac{ke^{-k(r-r0)}}{(e^{-k(r-r0)}+1)^2} = kA_\infty(r)(1 - A_\infty(r))$$

and thus, at steady state

$$J = \begin{bmatrix} -1 + wr(1-r) & -br(1-r) \\ \dfrac{1}{\tau}a(1-a) & -\dfrac{1}{\tau} \end{bmatrix}$$

We define $I_{1/2}$ as the level of drive for which there is a fixed point at $r = a = 0.5$, and thus

$$r = R_\infty(wr - ba + I)$$

$$0.5 = R_\infty\left(w(0.5) - b(0.5) + I_{1/2}\right)$$

$$I_{1/2} = R_\infty^{-1}(0.5) + 0.5(b - w)$$

$$I_{1/2} = x_0 - 0.5(w - b)$$

When there is sufficient recurrent excitation for UP/DOWN alternations (i.e. $w > w_0$, see derivation of $w_0$ below), $I_{1/2}$ gives the level of drive for equi-duration UP and DOWN states, for a given level of recurrence and adaptation strength (Figure S3).

If we define $I^* = I - I_{1/2}$ as the drive relative to $I_{1/2}$, the bifurcation diagram in $I^*$-$w$ parameter space (Supplemental Figure 4B) reveals that $I^* = 0$ (i.e. $I = I_{1/2}$) acts as an axis of symmetry of the effective I/O curve, with Excitable$_{\text{DOWN/UP}}$ regimes surrounding a bistable or oscillatory regime that has equi-duration UP/DOWN states at $I^* = 0$, depending on the values of $w, b$. Furthermore, transitions of the dynamic regime at the center of the I/O curve happen at $I^* = 0$. Supplemental Figure 4C shows the bifurcations at $I^* = 0$ with changing $w$ and fixed $b = 1$. As $w$ is increased, the fixed point at $r = 0.5$ loses stability with the appearance of oscillations in a Hopf bifurcation at $w = w_0$. With further increased values of $w$, two stable fixed points appear at high and low rate, marking the transition from oscillations to bistability in a pair of saddle node bifurcations at $w = w_\chi$. Finally, the pair of "inner" unstable fixed points coalesce in a pitchfork bifurcation at $w = w_{PF}$. Thus, the bifurcations at $I^* = 0$ reveal the parameter values at which qualitative changes in the I/O curve occur, between monotonic stable, oscillatory-centered, and bistable-centered I/O curves (Figure 3B).

To solve for the type of fixed point at $I = I_{1/2}$, we use the total input

$$X = wr - ba + I_{1/2}$$

$$X = w(r - 0.5) - b(a - 0.5) + x_0$$

For the fixed point at $[r^*\ a^*] = [0.5\ 0.5]$, $X = x_0$, giving

$$J = \begin{bmatrix} -1 + \frac{w}{4} & -\frac{b}{4} \\ \frac{k}{4\tau} & -\frac{1}{\tau} \end{bmatrix}$$

which we can use to obtain the conditions for $w_0$ and $w_{PF}$.

The condition for a Hopf bifurcation at $w_0$ is that $J$ has a pair of purely imaginary eigenvalues $\lambda_\pm = 0 \pm bi$ (Strogatz). $\lambda_\pm$ can be found by

$$\lambda_\pm = \frac{1}{2}\left(\mathrm{Tr} \pm \sqrt{\mathrm{Tr}^2 - 4\mathrm{Det}}\right)$$

where $\mathrm{Tr}$ and $\mathrm{Det}$ are the trace and determinant of $J$, respectively. The value for $w_0$ is found from the null-trace condition:

$$\mathrm{Tr} = 0$$

$$\left(-1 + \frac{w_0}{4}\right) + \frac{-1}{\tau} = 0$$

$$w_0 = 4\left(1 + \frac{1}{\tau}\right)$$

and minimal value for b follows from:

$$0 < \mathrm{Det}$$

$$0 < \left(-1 + \frac{w_0}{4}\right)\left(\frac{-1}{\tau}\right) - \left(\frac{b}{4}\right)\left(\frac{k}{4\tau}\right)$$

$$b > \frac{16}{\tau k}$$

The pitchfork bifurcation at $w_{PF}$ is the transition from 5 (3 unstable) to 3 (1 unstable) fixed points. $w_{PF}$ satisfies the condition that $J$ has a degenerate pair of eigenvalues $\lambda_\pm = 0$.

$$0 = \frac{1}{2}\left(\mathrm{Tr} \pm \sqrt{\mathrm{Tr}^2 - 4\mathrm{Det}}\right)$$

$$0 = \mathrm{Det}$$

$$0 = \left(-1 + \frac{w_{PF}}{4}\right)\left(\frac{-1}{\tau}\right) - \left(\frac{b}{4}\right)\left(\frac{k}{4\tau}\right)$$

$$w_{PF} = \frac{bk}{4} + 4$$

The degenerate pair of saddle node bifurcations at $w_{\chi}$, which separates oscillatory-centered and bistable-centered I/O curves, is determined numerically using XPP.

Together, this analysis reveals the shape of the I/O curve in the w-b parameter space. For low levels of recurrence, the I/O curve increases monotonically with a stable fixed point for each I-value and no UP/DOWN alternations are possible. At a critical value of recurrence, UP/DOWN alternations emerge at the axis of symmetry of the I/O curve. When adaptation is very weak, only bistability is possible and UP/DOWN alternations emerge in a cusp bifurcation as in Supplemental Figure 2C. With sufficient adaption UP/DOWN alternations emerge in Hopf bifurcation as in Supplemental Figure 3D. With sufficient recurrent excitation, the population will have a bistable-centered I/O curve if recurrence is stronger (Figure 3B, yellow) or an oscillatory-centered I/O curve if adaptation is stronger (Figure 3B, blue). Thus, the relative strength of recurrence and adaptation defines a spectrum from bistable-centered to oscillatory-centered response properties in the model.

*Physiological interpretation of model parameters*

Our model (Eqns 1,2) describes the mean activity of a neuronal population with positive feedback from recurrent excitation ($wr$), slow negative feedback from adaptation ($ba$), and a source of noisy drive ($I + \xi(t)$). We interpret drive in the model as the combination of external input and various other factors that drive cells toward spiking, including internal (for example, miniEPSPs) and modulatory influences (for example, increasing the excitability of cells would correspond to an increase in drive parameter in our model). The recurrence parameter, $w$, reflects the effective weight of excitatory synapses within the population. This self-excitation drives the population during the UP state. In the two-variable model, the saturating I/O relation, $R_{\infty}(input)$, imposes a maximum activity at $r(t) = 1$. However, the three-variable model reveals that fast negative feedback from inhibitory cells can dynamically stabilize the rate of excitatory population during the UP state.

Adaptation could encompass a variety of physiological processes that are activated by, and subsequently reduce, spiking at a slow timescale (~50ms-1s), such as slow voltage or calcium activated potassium currents, or sodium current inactivation [3,4]. For mathematical simplicity, we have reduced the effect of adaptive processes, to a single, saturating, mean field variable, $a(t)$, that negatively feeds back on population activity with strength $b$. While we have used a sigmoidal activation function for adaptation, this decision is not crucial for the dynamics described. With a linear activation function we would have been able to get similar results (albeit without the possibility for 5-Fixed point regimes). The choice of a sigmoid activation function was made for two reasons: 1) we found that sigmoid adaptation increases the robustness of the excitable regimes – it decreases the noise required for transitions out of stable fixed points and extends the parameter domain in which excitable alternations are seen. 2) Biologically, adaptation would be expected to saturate; for example, if adaptation were due to a voltage-gated ionic current. Previous studies have also modeled synaptic or slow divisive feedback to achieve the same goal [5,6], in some cases showing characteristic differences [5]. Further work to identify signatures of different sources of adaptive feedback could give insight to the implications and identification of biophysical substrates of adaptation in different physiological contexts.

With these idealizations, our model encompasses numerous previous models for UP/DOWN dynamics, from mean field to large scale spiking models. We have reduced the critical influences to a few key parameters that provide an intuitive understanding of a wide range of UP/DOWN alternation dynamics in neuronal populations.

*General insight to synchronized UP/DOWN dynamics*

Experiments in multiple physiological contexts have revealed that SWRs and UP/DOWN alternations are a locally-generated "default state" of hippocampal and neocortical tissue [7,8], ubiquitous under conditions of low neuromodulatory tone. Beyond NREM, they are observed during quiet wakefulness, [9,10] under anesthesia [11,12], and during *in vitro* slice or culture preparations [13,14] and in isolated tissue preprations [15,16]. Our model reveals why UP/DOWN alternations are so ubiquitous: they are seen in neural populations with sufficiently strong local recurrent excitation (>$w_0$) at comparatively low levels of drive ($\sim I_{1/2}$).

Synchronized UP/DOWN dynamics have been proposed to exist in a "multidimensional spectrum" [17]. Our model captures a subspace of this spectrum defined by two axes (Figure 7A). First, the magnitude of excitatory drive determines the degree of synchronization [2]. Increasing drive brings the population from a DOWN-dominated "more synchronized" regime with brief population bursts, to an UP-dominated "more asynchronous" regime with occasional DOWN states. The relative strength of recurrent excitation and adaptation determines the temporal dynamics between these extremes. When recurrence dominates, the system is characterized by bistability; when adaptation dominates, oscillations emerge. We note that this second axis also influences the "steepness" of UP/DOWN transitions, even in the adjacent excitable regimes, which we have not explored.

Different experimental conditions are associated with different regimes in the spectrum of UP/DOWN dynamics captured by our model. Other models have described oscillatory UP/DOWN dynamics in the spinal cord [18] and in cortical slices [19], whereas in cortical cultures $\text{Excitable}_{\text{DOWN}}$-like bursting behavior dominates [20]. Recently, bistable UP/DOWN dynamics were found to describe the activity of sensory cortex during urethane anesthesia [2,21]. During quiet wakefulness, cortical state varies following the level of arousal of the animal [22], but is unclear which of the regimes. Our model provides a framework by which one can predict how duration statistics should change with experimental manipulation of intrinsic or network properties, for example by varying levels of anesthesia or applying other pharmacological agents.

Extensive study has revealed multiple factors that can bring cortical tissue from the "default mode" of UP/DOWN dynamics to an activated state. In culture and slice preparations, increased levels of subcortical neuromodulators are able to 'wake up' the tissue, leading to the replacement of slow oscillations in the neocortex with asynchronous spiking [20] and SWRs in the hippocampus with theta-like oscillations [23]. Acetylcholine induces neocortical desynchronization and hippocampal theta during REM sleep[24], and acetylcholine, norepinephrine, or thalamic drive can give rise to cortical desynchronization during quiet wakefulness [25,26]. Each of these factors have effects that correspond to parameter changes that can transition our model from UP/DOWN dynamics to an asynchronous (tonic UP) state: 1) increasing excitability or external drive, $I$, 2) decreasing recurrence below the critical value for UP/DOWN alternations, $w < w_0$, or 3) decreasing the strength of adaptation, $b$, or 4) increasing recurrence such that the DOWN state loses stability. Similarly, the ascending neuromodulators increase excitability by depolarizing cortical pyramidal cells, decrease the effective weight of excitatory-excitatory synapses, and deactivate adaptive currents [27,28]. By these parallels, we are able to apply our model to interpret the putative mechanisms by which cortical desynchronization follows global and local levels of arousal.

*Description of the E-I-A model*

Our two-variable model captured significant features of UP/DOWN alternations and the relative roles of adaptation and recurrent excitation. However, firing rate in the UP state was described only as "active", limited by the saturating input-output function $R_\infty(input)$. One the other hand, neuronal spike rates during the UP state are generally low due to balanced excitation and inhibition. The e-i-a model represents the mean rate of an adapting excitatory and an inhibitory population,

$$\tau_e \dot{r_e} = -r_e + R_{e,\infty}\left(w_{ee} r_e - w_{ei} r_i - ba + I_e + \xi_e(t)\right) \qquad (3)$$

$$\tau_i \dot{r_i} = -r_i + R_{i,\infty}\left(w_{ie} r_e - w_{ii} r_i + I_i + \xi_i(t)\right) \qquad (4)$$

$$\tau_a \dot{a} = -a + A_\infty(r_e) \qquad (5)$$

with power law activation functions, as in Ahmadian and Milller (2013).

$$R_{e/i,\infty}(x) = k[x]_+{}^n$$

The activation function of adaptation, $A_\infty(r)$, is the same as in the r-a model, with parameters $r_0 = 2$, and $k = 3$. Unless otherwise specified, we've assumed for simplicity $w_{ee} = w_{ie} = w_e$ and $w_{ii} = 0$. However, the behaviors are robust to a range of weight values.
