## Supplementary material for "Excitable dynamics of NREM sleep: a unifying model for neocortex and hippocampus"

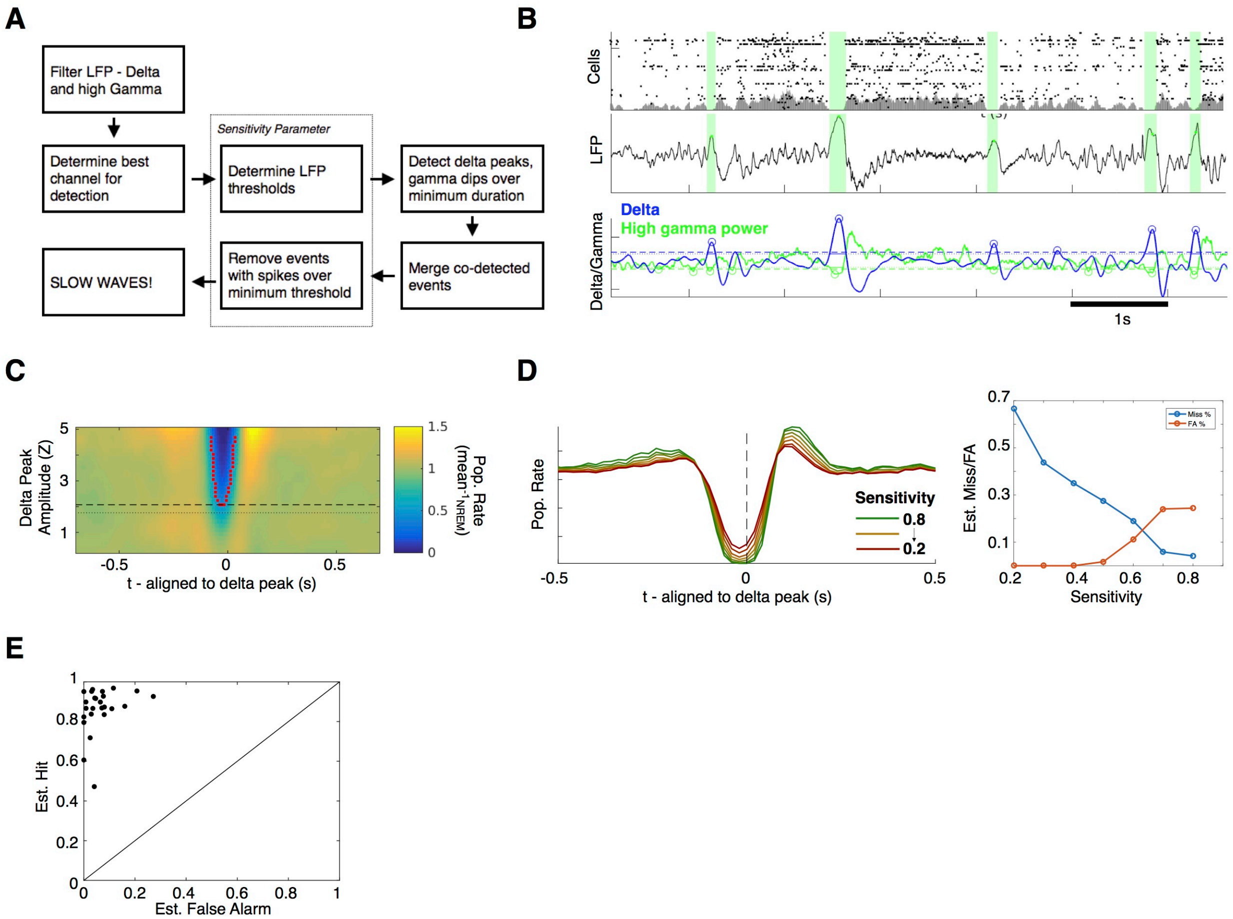


**Supplemental Figure 1 Slow wave detection. A:** The process for slow wave detection, as described in the Methods, is schematized. **B:** A 7s sample of neocortical data is shown with spike raster (top), LFP (middle), and filtered delta and high gamma power (bottom). Detection thresholds, as determined in panel C, are indicated as dashed (peak) and dotted (window) lines. Slow waves are detected as coincidence of threshold crossings of both signals. **C:** Threshold detection. Slow wave threshold was determined by calculating the population PETH around delta peaks, as a function of peak amplitude. Peak threshold (dashed line) was taken to be the lowest amplitude for which the PETH drops below a (mean-normalized) rate (given as the "sensitivity parameter"). Window threshold (dotted line) was taken to be the average value of the delta signal at which the PETH drops below the sensitivity parameter for any peak value (red dots). **D:** (Left) PETH around detected slow waves in the example recording in B,C for a range of sensitivity parameters. (Right) Estimated miss and false alarm % as a function of sensitivity parameter. Sensitivity = 0.6 was used for this study. **E:** Miss and false alarm % for each recording in the dataset.

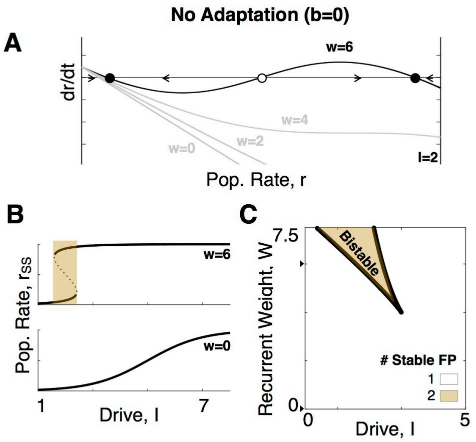


**Supplemental Figure 2: Recurrence enables UP/DOWN bistability. A:** Geometric analysis of the non-adapting (b=0) model illustrates the mechanism of bistability with sufficient recurrent excitation for fixed I=2. Curve indicates dr/dt for a given value of r. Circles indicate stable (filled) and unstable (empty) fixed points. **B:** Effective population **i**nput-output (I/O) curve, i.e. population rate steady state as a function of drive, for a population with low (w=0) and high (w=6) recurrence. **C:** I-W parameter space, bistable region appears in a cusp bifurcation at high recurrence and low input.

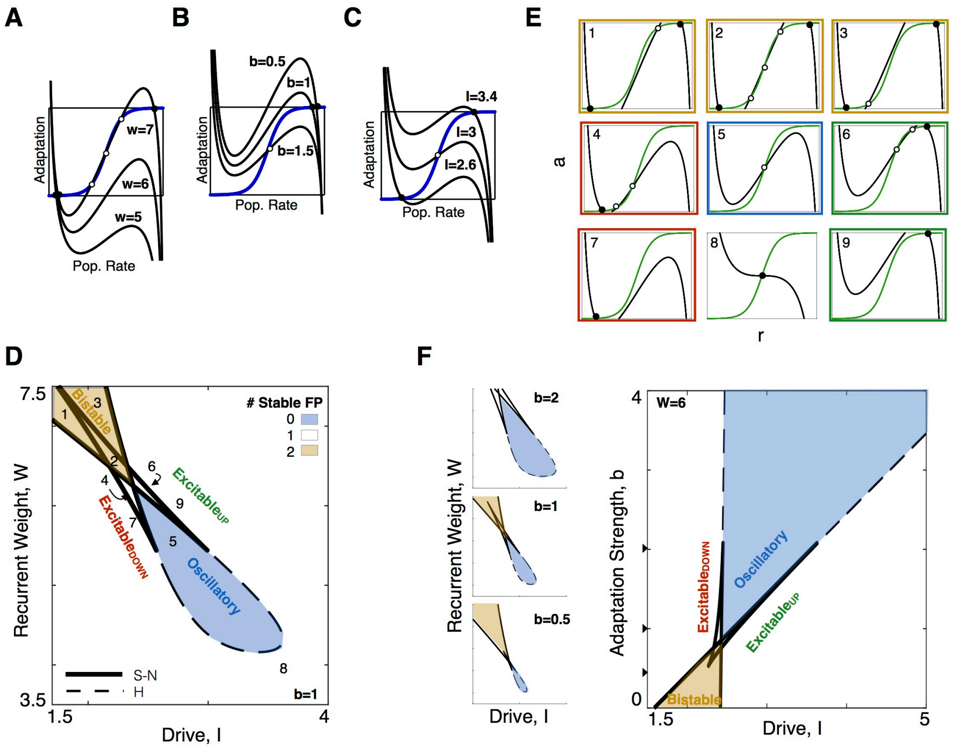


**Supplemental Figure 3 Parameter space of dynamical regimes of the r-a model.**
**A:** Effect of recurrence on the r-nullcline (b=1, I=2). **B:** Effect of adaptation strength, b, on r-nullcline (w=6, I=2.75). **C:** Effect of drive, I, on r-nullcline (w=5, b=1). **D:** I-W Bifurcation diagram reveals the "Butterfly Catastrophe" motif, yellow indicates bistability, blue indicates oscillations. Solid line, saddle-node (S-N) bifurcations; dashed line, Hopf (H) bifurcation. Numbers correspond to the phase planes in panel E. Note that while the 3-fixed point configuration in Figure 2Ciii/iv is limited to the small parameter domain of around number 4/6, excitable dynamics extend out to the 1-fixed point regions, as can be seen in Figure 3D. **E:** Representative phase plane from each domain of parameter space. Regime is determined by the location of stable fixed points. (Yellow: bistable, blue: oscillatory, red: Excitable$_{DOWN}$, green: Excitable$_{UP}$). **F:** The I-b parameter space. Increasing adaptation strength, b, increases the domain of the oscillatory regime. (Left) I-w parameter space for different values of b.

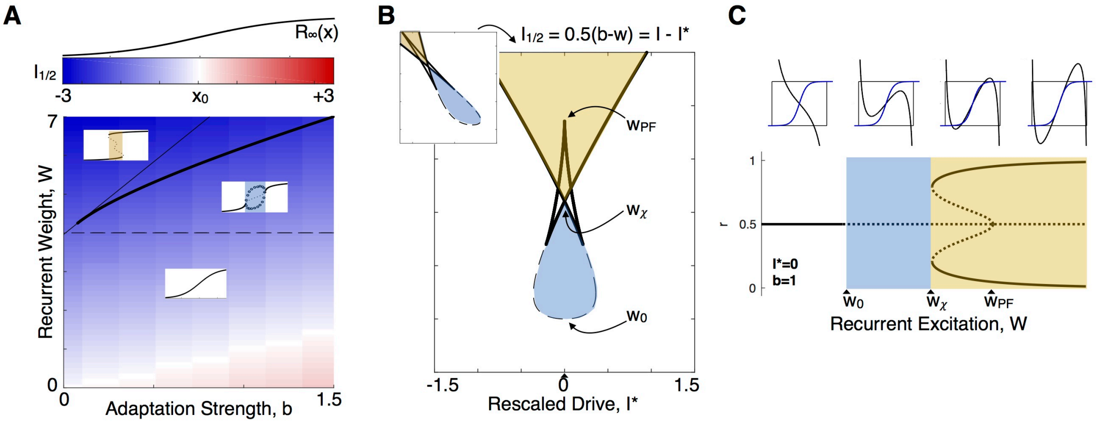


**Supplemental Figure 4 Dynamic regime at the I/O curve center region. A:** The level of drive drive for half-activation of the effective I/O curve ($I_{1/2}$, see Methods) varies as a function of w and b. Color indicates $I_{1/2}$ relative to $x_0$ (the midpoint of the unconnected I/O curve $R_\infty(x)$, shown as reference). Lines reflect bifurcations of the effective I/O curve, as shown in Figure 3A, revealing that the operating range for I shifts to lower values (deeper blue) when recurrence is sufficiently high for UP/DOWN alternations. **B:** I*-w parameter space. $I^*=I-I_{1/2}$ centers the parameter space around the I/O curve. **C:** Bifurcations in w, with $I^* = 0$ (i.e. at $I=I_{1/2}$). Oscillations appear in a hopf bifurcation at $w_0$, two stable (and two unstable) fixed points appear in a pair of saddle-node bifurcations at $w_\chi$, and the unstable fixed points coalesce with the center unstable fixed point in a pitchfork bifurcation at $w_{PF}$. Each of these bifurcations is shown in the w-b plane in Figure 3A.

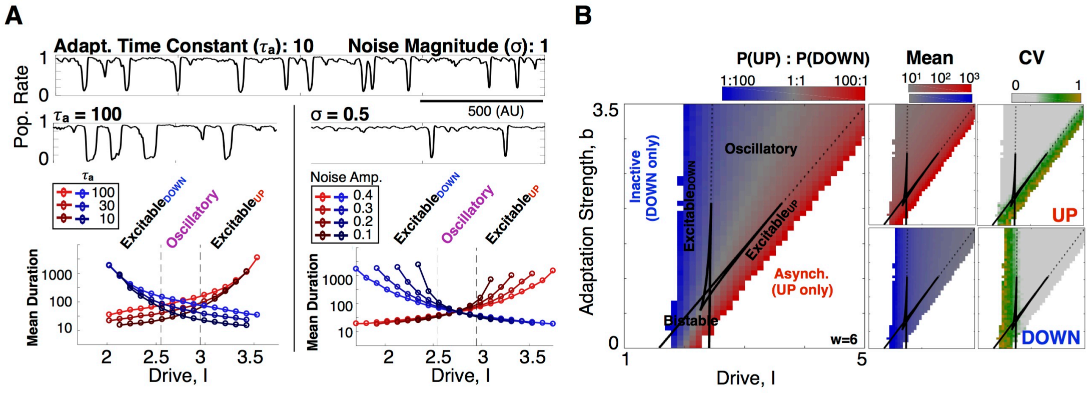


**Supplemental Figure 5 Effective I/O curve for a recurrent adapting population in the presence of noise. A:** Distinct effect of time scale of adaptation ($\tau_a$) and noise magnitude (σ) on stable and transient states. Increasing the time scale of adaptation increases the duration of transient states. Noise decreases the duration of stable states. **B:** Same as figure 3D, for I-b parameter space with fixed w.

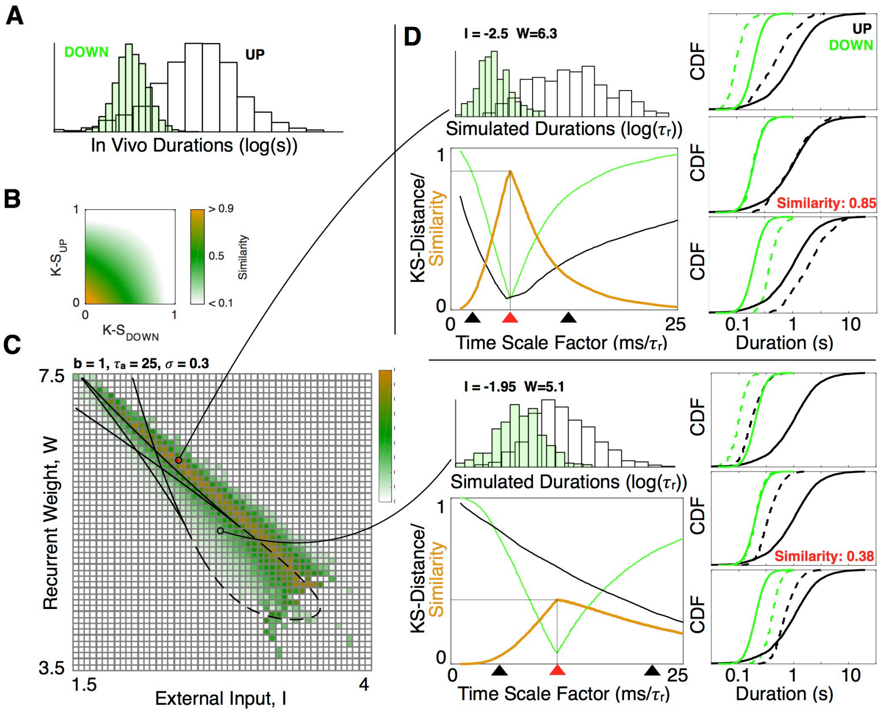


**Supplemental Figure 6 Duration distribution matching method. A:** UP/DOWN state duration distributions for an example recording. **B:** Similarity as a function of K-S distance between in vivo and simulated UP and DOWN state duration distributions. **C:** Map of similarity between the durations for the example recording in A and the r-a model with noise (Figure 3D). **D:** Calculation of similarity for two points in the I-w plane. For each point, KS distance between simulated and in vivo UP/DOWN state duration distributions is calculated at time scaling factors between 0.5 and 25ms/$\tau_r$. Simulated (dashed) and in vivo (solid) cumulative distributions are shown for three scaling factors (arrows) at each point. Similarity for the point is taken to be the similarity at the best time scale factor, re-dimensionalizing time to give the best match to the shapes and relative values of the UP and DOWN state duration distributions.

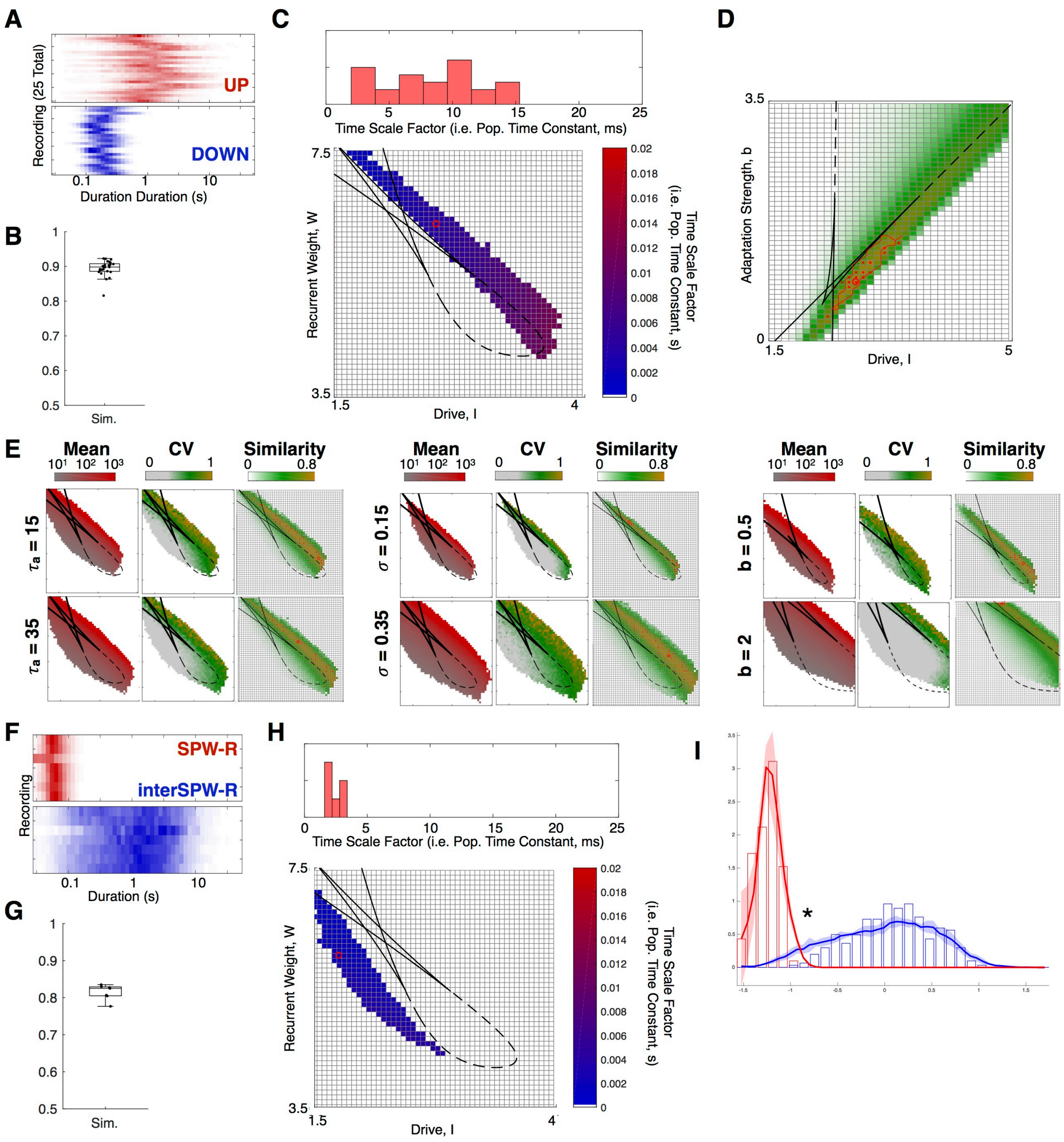


**Supplemental Figure 7** Distribution matching in Neocortex and Hippocampus. **A.** UP/DOWN state durations for all neocortical recordings in the dataset. **B:** Similarity metric for each neocortical recording the dataset. **C:** Histogram of time scale factors for all neocortical recordings in the dataset. (Bottom) mean time scale factor in I-w parameter space. **D:** Matching in the I-B parameter space. Same as figure 4B. **E:** Mean/CV of UP state durations and data-model similarity in the I-W plane with variation of the fixed parameters: time scale of adaptation, $\tau a$; magnitude of noise, $\sigma$, and adaptation strength, b. All other parameters same as in Figure 4B. **F-H:** Same as A-C, for hippocampus. **I:** Enlarged HPC duration distributions - model does not capture short duration interSWR periods.

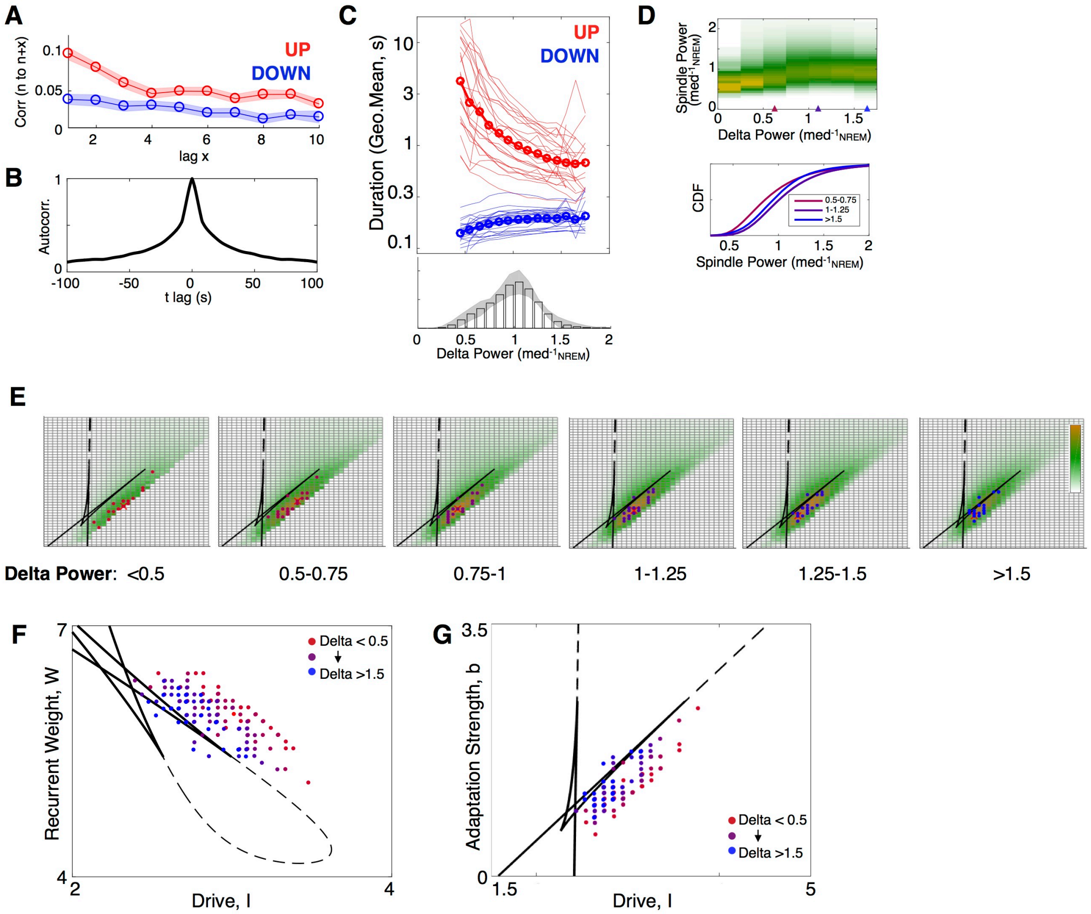


**Supplemental Figure 8 Slow variation in cortical state.** **A:** Correlation between durations of adjacent UP/DOWN states. **B:** Autocorrelation of delta power. **C:** Mean UP/DOWN state duration plotted vs delta power. Light lines: geometric mean duration for each recording. Dark lines: geometric mean duration over all recordings. **D:** High spindle power most common for intermediate delta power. Spindle power distribution as a function of delta power over all recordings. (Bottom) cumulative distribution of spindle power for low, intermediate, and high delta power. **E:** Same as Figure 6D but with matching to the I-b parameter space. **F,G:** Best matching parameter values in I-W and I-b parameter space for dwell time distributions in each recording, grouped by delta power (6 groups). Fitting procedure similar to Figure 4, with $\tau_r$ fixed at $\tau_r$=5ms.

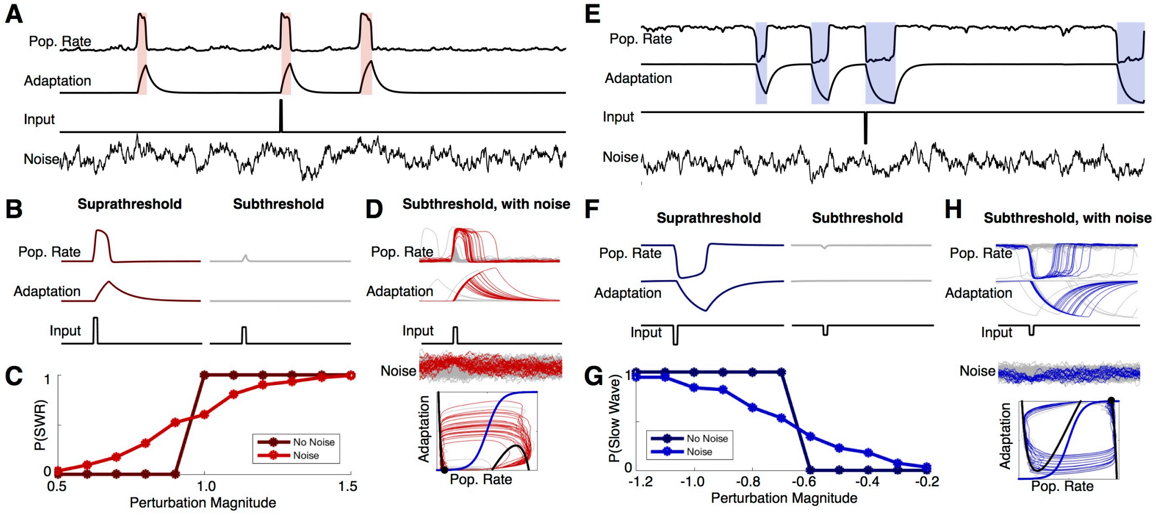


**Supplemental Figure 9. Perturbation-evoked Slow Waves/SWRs. A:** Simulated r-a model with best-matching parameters to hippocampal SWR dynamics, response to input perturbation ($I_{perturb}$ = 1.25). **B:** Suprathreshold ($I_{perturb}$ = 1.1) and subthreshold ($I_{perturb}$ = 0.7) perturbation of the noise-free model. **C:** Probability of perturbation-evoked SWR as a function of perturbation magnitude, with and without noise (magnitude: 0.2). **D:** Probabilistic SWR response to subthreshold perturbation in the presence of noise - colored trajectories are those in which an UP state was evoked. **E-H:** Same as A-D for the model with best-matching parameters to the cortical slow wave dynamics.

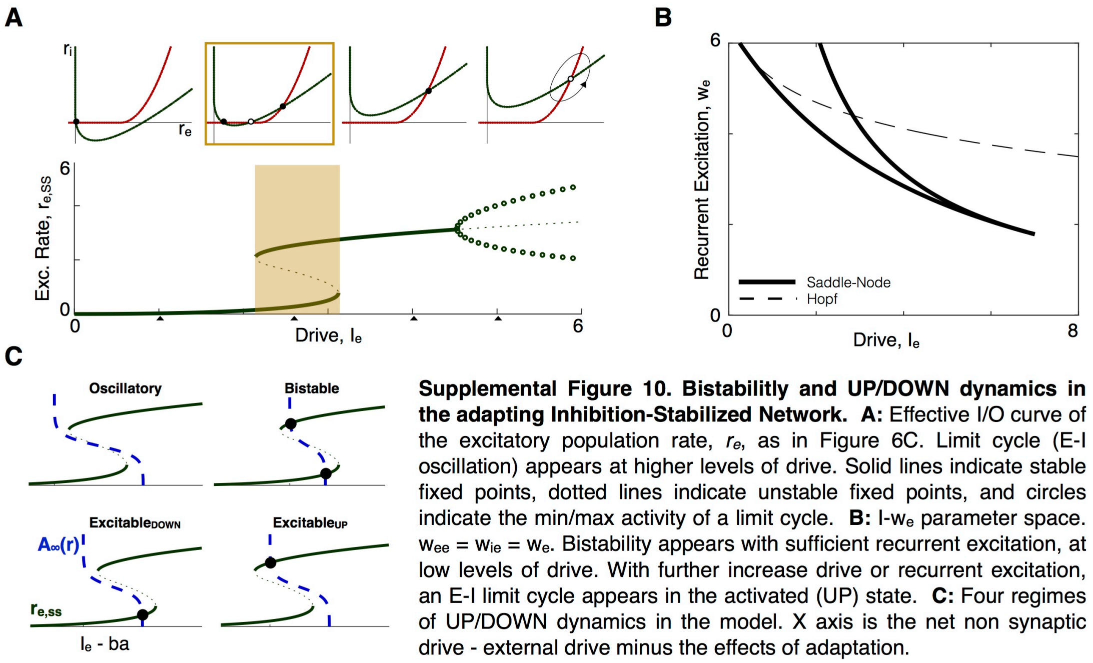


**Supplemental Figure 10. Bistabilitly and UP/DOWN dynamics in the adapting Inhibition-Stabilized Network. A:** Effective I/O curve of the excitatory population rate, $r_e$, as in Figure 6C. Limit cycle (E-I oscillation) appears at higher levels of drive. Solid lines indicate stable fixed points, dotted lines indicate unstable fixed points, and circles indicate the min/max activity of a limit cycle. **B:** I-$w_e$ parameter space. $w_{ee} = w_{ie} = w_e$. Bistability appears with sufficient recurrent excitation, at low levels of drive. With further increase drive or recurrent excitation, an E-I limit cycle appears in the activated (UP) state. **C:** Four regimes of UP/DOWN dynamics in the model. X axis is the net non synaptic drive - external drive minus the effects of adaptation.
